# MITOCHONDRIAL ANTIVIRAL PATHWAYS CONTROL ANTI-HIV RESPONSES AND ISCHEMIC STROKE OUTCOMES VIA THE RIG-1 SIGNALING AND INNATE IMMUNITY MECHANISMS

**DOI:** 10.1101/2024.06.07.598027

**Authors:** Silvia Torices, Thaidy Moreno, Sita Ramaswamy, Oandy Naranjo, Timea Teglas, Olivia M. Osborne, Minseon Park, Enze Sun, Michal Toborek

**Author notes:** **Corresponding Authors:** Silvia Torices, PhD and Michal Toborek, MD, PhD. Department of Biochemistry and Molecular Biology, University of Miami Miller School of Medicine, 528E Gautier Bldg. 1011 NW 15th Street, Miami, FL 11336.

## Abstract

Occludin (ocln) is one of the main regulatory cells of the blood-brain barrier (BBB). Ocln silencing resulted in alterations of the gene expression signatures of a variety of genes of the innate immunity system, including IFN-stimulated genes (ISGs) and the antiviral retinoic acid-inducible gene-1 (RIG-1) signaling pathway, which functions as a regulator of the cytoplasmic sensors upstream of the mitochondrial antiviral signaling protein (MAVS). Indeed, we observed dysfunctional mitochondrial bioenergetics, dynamics, and autophagy in our system. Alterations of mitochondrial bioenergetics and innate immune protection translated into worsened ischemic stroke outcomes in EcoHIV-infected ocln deficient mice. Overall, these results allow for a better understanding of the molecular mechanisms of viral infection in the brain and describe a previously unrecognized role of ocln as a key factor in the control of innate immune responses and mitochondrial dynamics, which affect cerebral vascular diseases such as ischemic stroke.

## INTRODUCTION

Following viral infection, the innate immune system employs pattern recognition receptors (PRRs), including transmembrane toll-like receptors (TLR) and RIG-I-like receptors (RLRs), to recognize extracellular and intracellular viral RNA, respectively 1–4. After binding with the ligand, the RLRs, such as the retinoic acid-inducible gene I (RIG-I) and the melanoma differentiation-associated protein 5 (MDA5), translocate to mitochondria to interact with the mitochondrial antiviral signaling protein (MAVS). MAVS activation results in the formation of the prion-like filament structure 5–7 and the activation of the downstream molecules TANK binding kinase-1 (TBK1) and interferon regulatory factor 3 (IRF3), which induce transcriptional expression of alpha/beta interferon (IFN-α/β) 5,8. Interestingly, MAVS activation is correlated with mitochondrial morphological changes and membrane disruption due to its pro-apoptotic function 9–13. Alterations in mitochondrial dynamics have been shown to affect the RLR signaling pathway, highlighting the role of mitochondria as a regulator of the immune response 14–17.

HIV-1 infection is often accompanied by a spike in IFN-α/β production 18–20. IFNs bind to their receptor complexes, which activate the signaling cascade through the Janus kinase signal transducer and activator of transcription (JAK/STAT) pathway that initiates the transcription of the IFN-stimulated genes (ISGs). ISGs play vital antiviral roles by inhibiting viral replication and production 21–24. Recently, our laboratory demonstrated that the ISG family 2’,5’-oligoadenylate synthetases (OAS) genes can regulate HIV-1 replication in BBB pericytes. However, the specific role of ISGs in HIV infection of this cell type remains largely unknown 25.

Human brain pericytes are key cells involved in the maintenance and regulation of the blood brain barrier (BBB) 26–29. Recent evidence indicates that BBB pericytes can be involved in immune responses, and that they can be infected by a variety of neurotropic viruses. Among them, brain pericytes can be infected by HIV-1, facilitating the release of the viral particles into the central nervous system (CNS) 25,30-36. Among several host factors involved in HIV infection, the protein occludin (ocln) has recently been shown to play a key role in regulating the extent of HIV-1 infection 25,31,35.

Ocln is a 65-kDa transmembrane protein formed by 522 amino acids (aa) 37. Although it is primarily known for its role in tight junction (TJ) assembly and function in barrier-generating cells, ocln is expressed as multiple spliced variants and has been shown to be a multifunctional protein whose barrier function is influenced by its phosphorylation status 38,39. It is formed by a long C-terminal cytoplasmic domain, four transmembrane domains, two extracellular loops, one intracellular loop, and a short N-terminal cytoplasmic domain 39,40. It has been reported that the N-terminus of ocln interacts with the WW domains of the E3 ubiquitin-protein ligase Itch, a target in MAVS for K48-linked polyubiquitination and degradation 41,42. Results from our laboratory have shown that ocln is present primarily in the cytoplasm in human brain pericytes 31,35, and may influence cellular metabolism by its function as an NADH oxidase and by modulating AMPK protein kinase activity 31,43. Being an important protein in preserving BBB integrity, ocln is a critical player in multiple brain disorders, such as ischemic stroke 44–48 as BBB damage is a common pathological feature in acute or chronic neurological disorders 49–52. Recently, altered ocln levels have been linked to psychiatric disorders such as bipolar disorder or autism spectrum disorder (ASD) 53,54.

In this study, we focus on the novel mechanisms regulating the impact of ocln on innate immune responses via modulating the IFN and the RLR signaling pathways both *in vivo* and *in vitro*. We demonstrate that ocln regulates HIV-1 infection by inducing the expression of antiviral ISGs and RIG-I, affecting MAVS activity, mitochondrial bioenergetics, and apoptosis in human brain pericytes. We also provide novel results that ocln deficiency worsens ischemic stroke outcomes in HIV-infected brains. Collectively, our findings establish ocln as a key regulator of the innate immune responses and HIV-1 infection. Moreover, our results point to ocln as a critical player in controlling cerebrovascular diseases such as ischemic stroke, providing a potential therapeutic target in these diseases.

## MATERIAL AND METHODS

### Cell Culture

Primary human brain vascular pericytes (ScienCell, Carlsbad, CA, USA, Cat#1200) corresponding to six various lot numbers were cultured in pericyte growth medium (ScienCell, Cat#1201). The medium was enriched with fetal bovine serum (FBS, 2%), 100 units/mL penicillin, 100 µg/mL streptomycin, and pericyte growth factors. Cells from passages 2 to 7 were utilized. Additionally, human embryonic kidney HEK293T/17 cells (ATCC, Manassas, VA, USA, Cat# CRL-11268) were grown in DMEM media (Thermo Fisher Scientific, Carlsbad, CA, USA, Cat#11995-065) supplemented with FBS 10% (ScienCell, Cat# 0500), 100 units/mL penicillin, and 100 ug/mL streptomycin (Thermo Fisher Scientific, Cat#15140-122). All cells used were maintained at 37°C with 5% CO_2_.

### HIV-1 infection and quantification

HIV-1 pNL4-3 strain was obtained from the NIH AIDS Reagent Program (Division of AIDS, NIAID, National Institutes of Health). EcoHIV/NDK, created by replacing the viral gp120 protein with gp80 from murine leukemia virus 101, was a gift from Dr. David Volsky’s laboratory (Icahn School of Medicine at Mt. Sinai, New York, NY). The pNL4-3 and EcoHIV/NDK plasmid amplification was accomplished using Stbl3 competent cells (Thermo Fisher Scientific, Cat# C737303). A total of 50 µg of proviral plasmid was transfected into 10^7^ HEK293T/17 cells for virus production using Lipofectamine 2000 (Thermo Fisher Scientific, Cat# 11668-027). The next day, cell media was replaced with fresh Opti-Mem (Thermo Fisher Scientific, Cat# 11058-021). After 48 hours of incubation, the supernatant was filtered using 0.45 µm-pore size filters (Millipore Sigma, Massachusetts, MA, USA, Cat# 430314) and concentrated using 50 kDa molecular weight exclusion columns (Millipore Sigma, Cat# UFC905024). Aliquots were stored at −80°C, and virus level production was assessed by quantification of p24 using the HIV-1 p24 Antigen ELISA 2.0 assay (Zeptometrix, Buffalo, NY, USA, Cat# 0801008). The levels of p24 were calculated in pg/ml. For *in vitro* HIV-1 infection, human brain pericytes were incubated with 60 ng/ml of HIV-1 p24. The cells were then extensively washed with PBS to remove the unbound virus, and fresh pericyte medium was added.

### Ocln deficient mice and EcoHIV infection

Ocln^+/−^ mice (Kumamoto University, Center for Animal Resources and Development, Kumamoto, Japan) were bred in the University of Miami animal facility, authenticated by genotyping, and assigned to the ocln^−/−^, ocln^+/−^, or ocln^+/+^ groups. Age-matched mice were used in all experiments. All animal experiments were consistent with the National Institutes of Health (NIH) guidelines, and all procedures were performed following the protocols approved by the University of Miami Institutional Animal Care and Use Committee (IACUC).

For *in vivo* EcoHIV infections, virus (1 µg of p24) was infused via the internal carotid artery as previously described 55. Briefly, the common carotid artery and its branching toward the internal and external carotid arteries were exposed. A cannula was inserted into the external carotid artery and EcoHIV was infused directly into the internal carotid artery. The animals were sacrificed one week after the surgery, and the tissue was harvested at the same time. Mouse EcoHIV infection rates were measured by q-PCR using the following primers and probe: NDKgag_F 5′-GAC ATA AGA CAG GGA CCA AAG G-3′; NDKgag_R 5′-CTG GGT TTG CAT TTT GGA CC-3′; NDKgag_Probe 5′-AAC TCT AAG AGC CGA GCA AGC TTC AC-3′. Normalization was performed based on HBB for DNA: MGBF 5′-CTG CCT CTG CTA TCA TGG GTA AT-3′, MGBR 5′-TCA CTG AGG CTG GCA AAG GT-3′; MGBP_Probe 5′-TTA ACG ATG GCC TGA ATC-3′. EcoHIV and mHBB standard curves were performed in parallel to evaluate copy numbers.

### Middle cerebral artery occlusion (MCAO)

An ischemic stroke was induced by the middle cerebral artery occlusion (MCAO) technique one week after mock or EcoHIV-infection. Briefly, middle cerebral artery blood flow was blocked by inserting a silicon-coated suture as previously described 87. After 60 minutes, the silicone suture was removed, and reperfusion was allowed for 24 h. The harvested brains were then sectioned using a 1 mm brain matrix and stained with 2,3,5-triphenyltetrazolium chloride (TTC) (Sigma, St. Louis, MO, Cat# T8877-10G). Stroke infarct volume was measured using Image J and corrected for edema using the following formula: Infarct size = [ipsilateral volume × infarct volume]/contralateral volume 87,102.

### Brain microvessel isolation

Briefly, mice were sacrificed, and the brains were immediately immersed in an ice-cold preservation solution (NaCl 5.98 g/L, KCl 0.35 g/L, KH_2_PO_4_ 0.16 g/L, MgSO_4_ 0.144 g/L, CaCl_2_ 0.30 g/L, HEPES 3.9 g/L, Glucose 1.8 g/L, NaHCO_3_ 2.10 g/L, and sodium pyruvate 0.11 g/L). Brains were homogenized using a Con-Torque Tissue Homogenizer (Eberbach, Van Buren Charter Township, MI, Cat# E2355.00). The resulting homogenized samples were filtered through a 300 μm nylon mesh filter (Fisher Scientific, Cat# NC1480938), gently agitated with 26% dextran, and centrifuged at 5800g for 30 min at 4 °C. The pellet was resuspended in an ice-cold preservation solution, passed through a 120 μm nylon mesh filter (Merck, St. Louis, MO, Cat# NY2H09000), and centrifuged at 1500g for 10 min at 4 °C. The pellet was resuspended in adequate RNA extraction buffer, and RNA was isolated as previously explained.

### Neurodeficit testing

Neurological outcome after ischemic stroke was measured using a scale adapted from Cuomo et al 56. This scale allows for the evaluation of motor function and behavior recovery. To avoid variability in the experiments, the investigators were blinded, and all tests were performed in the same dedicated laboratory and at the same time of the day.

### RNA libraries and sequencing

For the RNAseq analysis, five independent replicates were prepared separately for non-specific small interfering RNA (siRNA) controls and ocln siRNA. Total RNA was isolated using the iPrep PureLink Total RNA kit (Thermo Fisher Scientific, Cat#12183018A) according to the manufacturer’s instructions. RNA quantitation and quality were assessed using the Qubit RNA HS Assay Kit (Thermo Fisher Scientific, Cat#Q32855) and the Agilent RNA 6000 Pico Kit (Agilent, Santa Clara, CA, USA, Cat#5067-1513). RNA-Seq libraries were prepared using the TruSeq mRNA Library Prep Kit (Illumina, Madison, WI, USA). To evaluate fragment libraries, an Agilent 2100 Bioanalyzer, a High Sensitivity DNA Kit (Agilent, Cat#5067-4626), and a qPCR-based KAPA library quantification kit (KAPA Biosystems, Wilmington, MA, USA, Cat#KK4600) were employed. Libraries were pooled into a 20 pM final concentration and submitted to the sequencing reaction using the MiSeq Reagent Kit (Illumina) and the MiSeq Sequencing System (Illumina).

### RNA-seq data analysis

Illumina reads were trimmed to remove low-quality bases from the ends and aligned using the CLC Genomics Workbench platform (Qiagen, Hilden, Germany, EU). The total gene hit counts and reads per kilobase million (RPKM) were calculated using the same platform. To determine differential gene expression associated with ocln silencing, comparisons were performed between the five replicates from cells transfected with ocln siRNA and the five replicates from controls transfected with non-specific siRNA. Differential gene expression (DEG) analysis was performed using the DESeq2 R package 103. An adjusted p-value of 0.05 and an absolute log2 fold change of 1 were considered cut-offs to generate the differentially expressed gene list 104. Computed z-scores of significant genes are represented in the heatmap. The heatmap was plotted using the pheatmap R package version 1.0.12. (https://CRAN.R-project.org/package=pheatmap).

The DEG list was employed to perform pathway enrichment and interaction network analysis using the g:GOSt tool from the g:Profiler web (https://biit.cs.ut.ee/gprofiler/). Only annotated genes were used to define background genes for comparison; multiple testing corrections were performed with g:SCS (p-value threshold = 0.05) for reducing significance scores as g:SCS takes into account the unevenly distributed structure of functionally annotated gene sets. In order to represent statistically significant related pathways, the pathway definition file and the output results from the g:Profiler tool were used as input in Cytoscape 105 (http://www.cytoscape.org/) and Enrichment Map 106 plugging (http://www.baderlab.org/Software/EnrichmentMap) to create the network visualization of pathways on Figure 1, laying out and clustering automatically by group of similar pathways into major pathways and reactions, following a previously published protocol 107. Reactome pathway terms were visualized as a chord plot using GOplot R 108.

**Figure 1.**
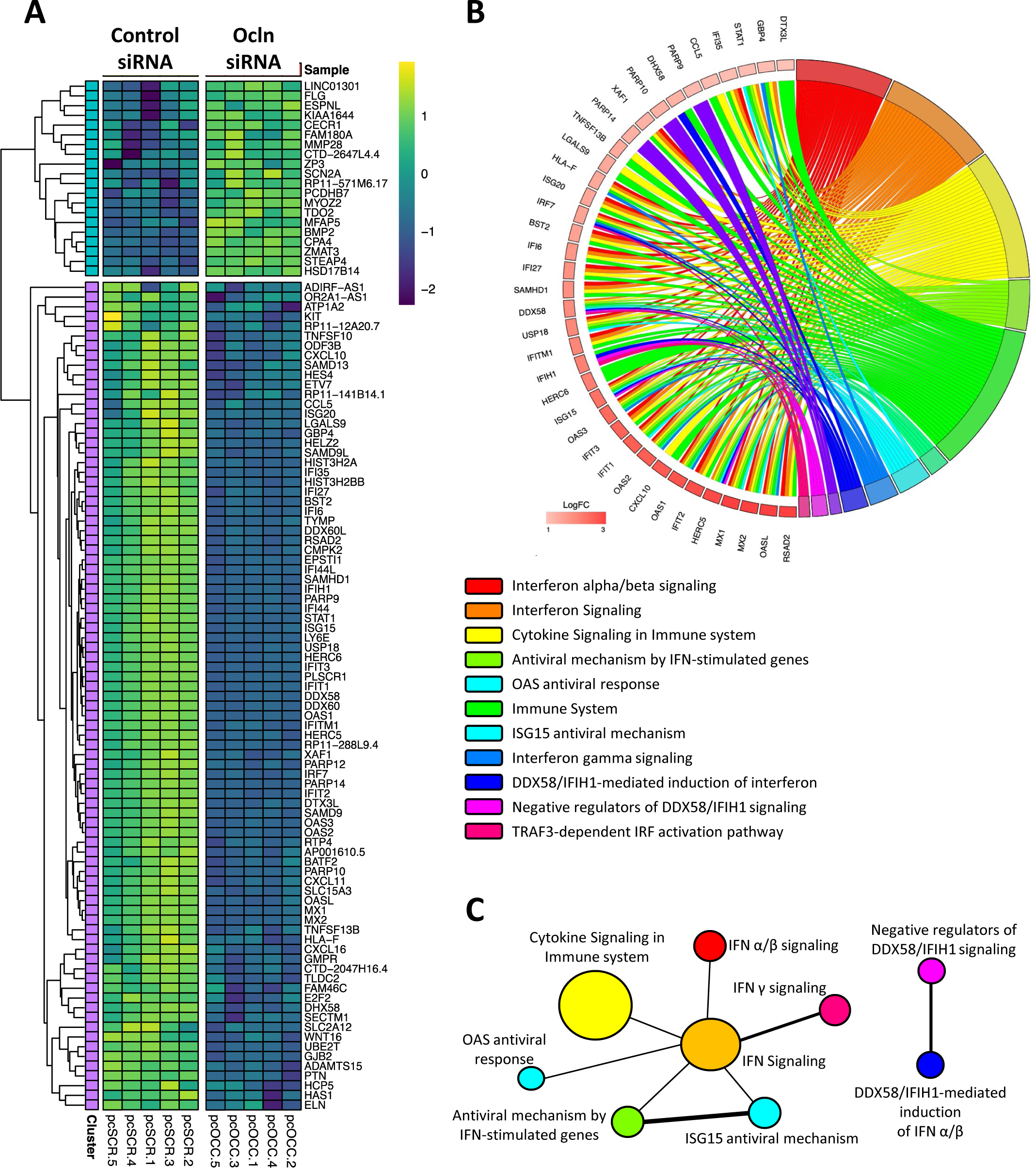
Ocln silencing displays distinct RNA-seq transcriptome signatures. **(A)** Heatmap of RNA-Seq expression z-scores computed for genes that are differentially expressed (*p* < 0.05, |log 2 (foldchange)| > 1) between the five replicates of ocln siRNA samples and the five replicates from the negative siRNA controls. **(B)** Chord diagram showing enriched Reactome clusters for the differentially expressed genes. Genes contributing to their respective enrichment are shown on the left, and enriched Reactome clusters are shown on the right. LogFC differentially expressed genes are displayed in shades of red. Each Reactome term is represented by one colored line. **(C)** The enrichment map is summarized by collapsing node clusters shown as circles (nodes), connected with lines (edges) of the Reactome pathways sharing genes. The network was manually adjusted to reduce node and label overlap.

### Real-Time PCR

To assess individual gene expression, mRNA isolation from human brain pericytes was performed using the Rneasy mini Kit (Qiagen, Cat# 74104) according to the manufacturer’s instructions. Genomic DNA from the mouse brain and spleen was isolated with the QIAmp DNA Mini Kit (Qiagen, Cat# 51304), mRNA from the plasma was isolated using the QIAamp MinElute Virus Spin Kit (Qiagen, Cat# 57704), and mRNA from brain microvessels was isolated using the QIAamp Lipid Tissue Mini Kit (Qiagen, Cat# 74804). Total RNA was quantified using Nanodrop 2000 (Thermo Fisher Scientific). A total of 100 ng of RNA was used in each reaction. Reverse transcription and qPCR reactions were performed at the same time by using the qScript XLT 1-Step RT-qPCR Tough Mix (Quantabio, Beverly, MA, USA, Cat# 89236-676). Gene amplification was achieved by employing TaqMan Gene Expression Assays and the following primers (all from Thermo Fisher Scientific): human-ISG15:Hs01921425_s1; human-IFIT1: Hs03027069_s1; human-MX1:Hs00895608_m1; human-MX2: Hs01550814_m1; human-HERC5:Hs00180943_m1; human-IRF7: Hs01287244_cn; human-IRF3: Hs01510490_cn; human-RIG-1: Hs01061433_m1; human-USP18:Hs00276441_m1; human-IFIH1: Hs00223420_m1; human-TRAF3: Hs00936781_m1; human-MAVS: Hs00920075_m1; human-MFN2: Hs00208382_m1; human-OPA1: Hs01047019_m1; human-FIS1: Hs00211420_m1; human-MIFF: Hs00697394_g1; human-PPARGC1α: Hs00173304_m1; human-CALCOCO2: Hs00977443_m1; human-P62: Hs01061917_g1; human-TJP1/ZO-1: Hs01551867_m1; human-OCLN: Hs05465837_g1; human-BCL2: Hs00608023_m1; human-BAX: Hs00180269_m1 and mouse-INFα5: Mm00833976; mouse-INFβ1:Mm00439552; mouse-INFɣ: Mm01168134; mouse-ISG15: Mm01705338; mouse-IFIT1: Mm07295796; mouse-RIG-1: Mm00487934; mouse-IFIH1: Mm00459183; mouse-FIS1: Mm00481580; mouse-BAX: Mm00432051; mouse-MFF: Mm00030720_cn; mouse-OPA1: Mm00451274_cn; mouse-BCL2: Mm00529770_cn; mouse-MAVS: Mm00523170_m1; mouse-TRAF3: Mm00385377_cn; and mouse-MFN2: Mm00500120.

GAPDH and β-actin mRNA levels were evaluated for sample normalization. Quantitative real-time PCR was performed using the Applied Biosystems 7500 system (Applied Biosystems, Foster City, CA). Gene expression changes were calculated by the ΔCt method, with Ct representing the cycle number at threshold. The desired PCR product specificity was determined based on melting curve analysis.

### Gene silencing and overexpression

For gene silencing, pericytes were transfected with the following human siRNAs (all from OriGene, Rockville, MD, USA): 3 unique 27-mer occludin-siRNAs (SR303274), 3 unique 27-mer TJP1-siRNAs (SR322042), 3 unique 27-mer ISG15-siRNAs (SR322849), 3 unique 27-mer Mx1-siRNAs (SR303015), 3 unique 27-mer Mx2-siRNAs (SR303016), or 3 unique 27-mer RIG-1-s RNAs (SR308383). Trilencer-27-universal scrambled (SCR) silencer siRNA duplex (OriGene, Cat# SR30004) was used as a negative control. For ocln overexpression, cells were transfected with the PCMV3-OCLN plasmid (Sino Biological, Wayne, PA, Cat# HG15134-UT) or PCMV3 (Sino Biological, Cat# CV011) as a negative control. A total of 0.5 µg per 10^6^ of siRNA and 1 µg per 10^6^ of DNA were transfected by Amaxa Nucleofector Technology using the Basic Nucleofector Kit (Lonza, Switzerland, EU, Cat# VPI-1001). After 18 h, the cells were washed with PBS and allowed to grow in pericyte medium.

### Immunoblotting

Following transfections, pericytes were washed with PBS and lysed with 100 µl/well of RIPA buffer containing protease inhibitors (Santa Cruz Biotechnology, Dallas, TX, USA, Cat# sc-24948a). Protein levels were quantified by the BCA Protein Assay Kit (Thermo Fisher Scientific, Cat# 23223). Samples were split and electrotransferred using 4-20% Mini-PROTEAN TGX Stain-Free Protein Gels and the PVDF membrane Trans-Blot Turbo Transfer System (Bio-Rad Laboratories, Hercules, CA, USA, Cat# 170-4159). Blots were blocked with bovine serum album (BSA) at 5% in TBS-0.05% Tween20 for 1 h. Next, they were incubated overnight at 4°C with the following primary antibodies (all at 1:1000 in 5% BSA-TBS): anti-occludin (Thermo Fisher Scientific, Cat# 71-1500), anti-ISG15 (Thermo Fisher Scientist, Cat# 703131), anti-MX1 (Cell Signaling, Danvers, MA, USA, Cat# 37849), anti-MX2 (Cell Signaling, Cat# 43924), anti-IFIT1 (Abcam, Cambridge, UK, Cat# ab70023), anti-USP18 (Thermo Fisher Scientific, Cat# PA5-110555), anti-HERC5 (Thermo Fisher Scientific, Cat# PA5-114382), anti-DRP1 (Cell Signaling, Cat#14647), D-RP1 (616) (Cell Signaling, Cat# 44945), anti-RIG-1 (Thermo Fisher Scientific, Cat# 700366), anti-MDA5 (Abcam, Cat# ab283311), anti-MAVS (Thermo Fisher Scientific, Cat# 14341-1-AP), anti-TRAF3 (Abcam, Cat# ab239357), anti-IRF3 (Thermo Fisher Scientific, Cat# 703682), anti-p-IRF3 (Ser 396) (Thermo Fisher Scientific, Cat# B5-3195R), anti-TBK (Cell Signaling, Cat# 51872S), anti-p-TBK (Ser 172) (Cell Signaling, Cat# 5483S), anti-FIS1 (Abcam, Cat# ab156865), anti-MFN2 (Cell Signaling, Cat# 11925S), anti-OPA1 (Cell Signaling, Cat# 80471S), anti-MFF (Cell Signaling, Cat# 845805), anti-TOMM20 (Thermo Fisher Scientific, Cat# PA5-52843), anti-LC3BI/II (Cell Signaling, Cat# 4108), anti-P62 (Cell Signaling, Cat# 16177), anti-NDP52 (Cell Signaling, Cat# 60732), anti-PGC1α (Cell Signaling, Cat# 2178), anti-BAX (Cell Signaling, Cat# 2772), anti-BCL2 (Cell Signaling, Cat# 3498). Samples were then washed with TBS-0.05% Tween20 three times for 5 min and incubated for 1 h with the secondary antibodies, all from LI-COR (Lincoln, NE, USA, Cat# 926-32210, Cat# 926-68070, Cat# 926-32211, Cat# 926-68071) at 1:20000 in 5% BSA-TBS. Anti-GAPDH antibody (1:20000, Novus Biologicals, Woburn, MA, USA, Cat# NB600–502FR or Cat# NB600-5021R) was used to normalize protein levels. Immunoblots were washed with TBS-0.05% Tween20 three times and analyzed using the Licor CLX imaging system and the Image Studio 4.0 software (LI-COR).

### Immunostaining

Human brain pericytes were cultured on a glass coverslip coated with 10 mg/mL of poly-L-lysine (Millipore Sigma, Cat# P8920). Cells were fixed and permeabilized with 4% paraformaldehyde for 15 min, followed by PBS containing 0.2% Triton X-100 for 15 min. Afterwards, cells were washed with PBS and blocked for 1 h with 3% BSA in TBS. The coverslips were then incubated with anti-TOM20 antibody (1:100 in 3% BSA in TBS; Thermo Fisher Scientific, Cat# PA5-52843) or anti-occludin (1:100 in 3% BSA in TBS; Thermo Fisher Scientific, Cat# 71-1500) overnight at 4°C in a humidified atmosphere. The next day, coverslips were washed three times with PBS and incubated for 1 h with Alexa Fluor 488-(Thermo Fisher Scientific, Cat# A12379) or 647-(Thermo Fisher Scientific, Cat# A31571) secondary antibodies diluted at 1:400 in 3% BSA in TBS. The coverslips were washed with PBS and mounted with Vectashield Antifade Mounting Medium with DAPI to visualize the nuclei (Vector Laboratories, Burlingame, CA, USA, Cat# H-1500). All images were obtained with the Olympus FLUOVIEW 1200 Laser Scanning Confocal Microscope (Olympus, Center Valley, PA, USA). For analysis of mitochondrial footprint and mean branch length, the Stuart Lab Mitochondrial Network Analysis 66 (v. 3.0.1, https://github.com/StuartLab) plugin was used 109.

### Mitochondria functional assessment

Mitochondrial respiration capacity was assessed by the Seahorse Cell Mito Stress Test Assay (Agilent, Cat# 103015-100) using a Seahorse XF24 Analyzer (Agilent Technologies, Inc.). A total of 1 × 10^5^ cells per well were seeded on a 24-well Xfe96 Cell culture Microplate (Agilent, Cat# 100777-004) coated with poly-L-lysine (Millipore Sigma, Cat# P8920) and grown overnight in the cell incubator. Oxygen consumption rate (OCR) and extracellular acidification rate (ECAR) were measured at basal levels and after adding the following inhibitors: 1.5 µM oligomycin (ATP synthase inhibitor), 1 µM FCCP (mitochondrial uncoupler), and 0.5 µM rotenone (complex I inhibitor)/antimycin A (complex III inhibitor). Results were analyzed using the Report Generator software, and values were normalized to total protein per well. Each data point reflected the average of six measurements.

### Measurement of ROS production

Mitochondrial ROS production was measured using the MitoSox Red mitochondrial superoxide indicator (Thermo Fisher Scientific, Cat # M36008), following the manufacturer’s instructions. Briefly, after ocln transfection, cells were seeded at a density of 1.5×10^5^ cells/well on 48-well plates. The next day, cells were incubated with 5 μM MitoSOX reagent in Ca/Mg-free HBSS buffer for 15 min in the dark, followed by washing three times with DPBS. Fluorescence was recorded using the SpectraMax iD3Reader (Molecular Devices, USA) at excitation and emission wavelengths of 510 and 580 nm, respectively.

### Statistical Analysis

Except for DESeq2 differential gene expression, statistical analyses were performed using GraphPad Prism Software version 6.0 (La Jolla, CA, USA). Statistical significance was determined by one-or two-way ANOVA followed by Tukey’s multiple comparisons test or Student’s *t* test. The significance level was set at *p*<0.05 (\*\*\*\**p*<0.0001, \*\*\**p*=0.0002, \*\**p*=0.003, \**p*<0.0449).

## RESULTS

### Cellular ocln levels control the expression of antiviral ISG genes

To evaluate the involvement of ocln in cellular functions of BBB pericytes, cells were transfected with ocln targeted siRNA for 48 h. Differential gene expression in ocln-silenced cells was analyzed via high-throughput RNA sequencing (RNA-seq) and compared to pericytes transfected with control siRNA. A total of 14415 genes were analyzed in the obtained dataset. The heatmap and volcano plot (**Fig. 1A and Suppl. Fig. 1A**, respectively) visualized 105 downregulated or overexpressed genes with a fold change > 1 in ocln-silenced pericytes as compared to controls. The heatmap of RNA-seq expression z-scores clearly identified two major clusters of downregulated and upregulated genes. The number of downregulated genes (85 genes) markedly exceeded the number of upregulated genes (20 genes) in pericytes with silenced ocln gene.

We then performed gene set enrichment analysis (GSEA) to identify pathways affected by ocln silencing. The chord diagram summarizes the enriched reactome pathways for target genes differentially regulated by ocln silencing (**Fig. 1B**). Genes contributing to their respective enrichment are shown on the left, and enriched reactome clusters are shown on the right. LogFC of differentially expressed genes are displayed in shades of red. The following eleven pathways were altered after ocln silencing when the analysis was restricted to the genes with a fold change higher or equal to 1: IFN-α/β signaling, IFN signaling, cytokine signaling in immune system, antiviral mechanism by IFN-stimulated genes (ISG), OAS antiviral response, immune system, ISG15 antiviral mechanism, INFɣ signaling, DDX58/IFIH1-mediated induction of IFN-α/β, negative regulators of DDX58/IFIH1 signaling, and TRAF3-dependent IRF activation pathway (**Fig. 1B and Suppl. Fig. 1B**). The *p*-values for these reactome-enriched pathways and the specific genes affected in each pathway are summarized in **Suppl. Fig. 1B and Suppl. Table 1**. **Fig. 1C** illustrates the overlaps among enriched pathways by collapsing node clusters shown as circles (nodes), connected with lines (edges) of the reactome pathway sharing genes.

In this initial series of experiments, we identified that silencing cellular ocln levels affected the expression of IFN-related genes and pathways. In order to establish the specificity of these responses, parallel experiments were performed on pericytes with silenced ZO-1, another TJ protein. Pericytes were transfected with ocln siRNA, PCMV3-OCLN vector for ocln overexpression, or ZO-1 siRNA, and the expression levels of selected ISG genes were analyzed by q-PCR and immunoblotting. **Suppl Figs. 2A-B** illustrate the effectiveness of the employed manipulation on ocln and ZO-1 expression, respectively. The silencing of ocln resulted in a decrease in mRNA expression by 45%; however, ocln overexpression increased ocln level by ∼4000 times at the mRNA level and by 250% at the protein level. In addition, ZO-1 silencing decreased ZO-1 mRNA levels by 50%.

Ocln silencing resulted in significantly diminished mRNA expression of the genes of the ISG pathway (**Figs 2A-H**), such as ISG15, IFIT1, MX1, MX2, USP18 and HERC5 when compared with control siRNA (**Figs 2A, C, E-H**). In contrast, no changes were found in the expression of these genes following ZO-1 silencing, except for a reduction of MX2 levels (**Figs. 2F**), illustrating the specificity of responses. Moreover, ocln overexpression resulted in a significant increase in mRNA levels of ISG15, IFIT1, MX1, MX2, USP18, and HERC5 (**Figs. 2A, C, E-H**). Ocln overexpression also elevated protein levels of ISG15, MX1 and MX2, without changing IFIT1, USP18, or HERC5 protein expression **(Figs. 2A, C, E-H).** We also evaluated the IGS15 and IFIT1 mRNA expression in cerebral microvessels isolated from the brains of ocln^+/+^ (wild type controls), ocln^+/−^, and ocln^−/−^ mice. ISG15 mRNA levels were significantly lower in microvessels isolated ocln-deficient mice as compared to controls with normal ocln expression. However, no changes were found in IFIT1 expression levels (**Figs. 2B, D**).

**Figure 2.**
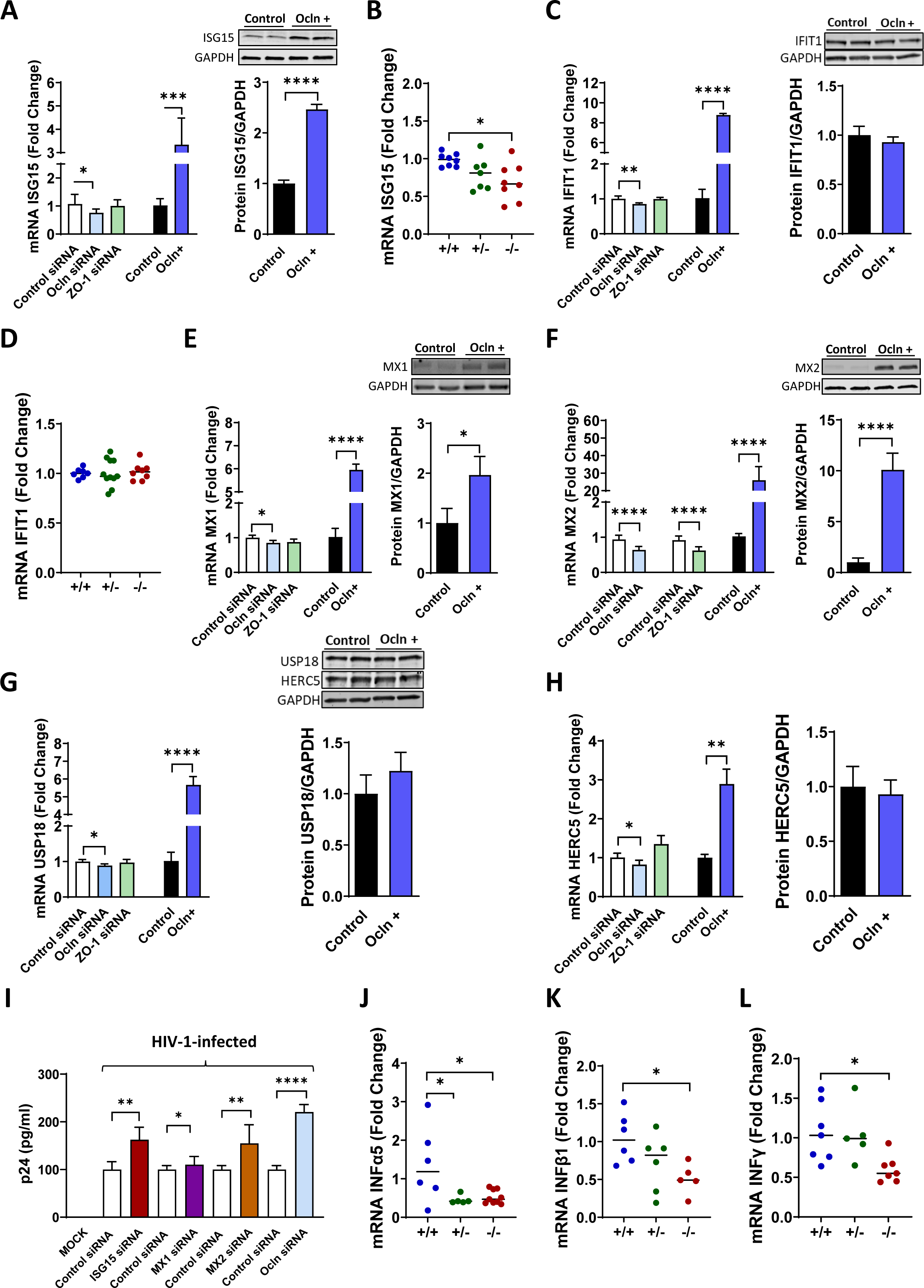
Ocln regulates HIV-1 infection through the ISG antiviral gene expression. Pericytes were transfected with ocln-siRNA, ZO-1 siRNA, or PCMV3-OCLN vector, and the expression of ISG15 **(A)**, IFIT1 **(C)**, MX1 **(E)**, MX2 **(F)**, USP18 **(G)**, and HERC5 **(H)** was evaluated by q-PCR and Western Blotting. Pericytes were transfected with ISG15-siRNA, MX1-siRNA, MX2-siRNA, and ocln-siRNA. Next, cells were either Mock-infected or infected with 60 ng/ml HIV-1 p24 for 12 hours. p24 levels were analyzed in cell culture media by ELISA as a marker of HIV infection **(I)**. n= 4-8 per group. RNA was extracted from isolated microvessels of age- and sex-matched ocln^−/−^, ocln^+/−^, and ocln^+/+^ mice (*n*= 6-8 animals per group). ISG15 **(B)**, IFIT1 **(D)**, IFNα5 **(J)**, IFNβ1 **(K)**, and IFNɣ **(L)** mRNA levels were analyzed by q-PCR. Graphs indicate the mean ± SD from three independent experiments. Individual animal data points are marked by blue (ocln^+/+^), green (ocln^+/−^), and red (ocln^−/−^) dots. ****p < 0.0001, \*\*\**p* = 0.0002, **p = 0.003, *p < 0.0449.

Given the importance of the ISG15 pathway in immune responses, we next investigated whether their altered expression, independent of ocln silencing, can influence HIV-1 infection of human brain pericytes. Cells were transfected with siRNA specific for ISG15, MX1, MX2, or ocln (positive control) and were either Mock-infected or infected with 60 ng/ml of HIV-1 for 12 h. **Suppl. Fig. 2C** represents the efficiency of gene silencing, which was ∼50% for each gene. The next day, cultures were washed to remove the unbound virus, and fresh medium was added. The levels of p24 antigen as a marker of HIV-1 infection were analyzed 48 h post-infection in the cell culture media by ELISA. Downregulation of ISG15 and MX2 resulted in enhanced HIV-1 replication. As expected, ocln silencing resulted in a comparable elevation of p24 levels. In contrast, no changes in p24 levels were detected after MX1 silencing (**Fig. 2I**).

This series of experiments was completed by evaluation of the INF genes in ocln-deficient mice. The INFα5, INFβ1, and INFɣ mRNA levels were significantly lower in microvessels isolated from ocln-deficient mice when compared to ocln control mice (**Figs. 2J-L**), confirming the critical role of ocln in regulation of interferon signaling and innate immunity responses.

### Ocln deficiency and EcoHIV infection worsen ischemic stroke outcomes

Reports from our laboratory 25,31,33,35,36, including data presented in **Fig 2I**, have shown that ocln can regulate the extent of HIV-1 infection in various cell types, including brain pericytes. However, no studies have been performed *in vivo* demonstrating the influence of ocln on HIV replication in the brain. Therefore, we evaluated the importance of ocln in host defense against HIV infection by employing ocln-deficient mice. Briefly, ocln^−/−^, ocln^+/−^, and ocln^+/+^ mice were infected with EcoHIV by infusion via the internal carotid artery for one week 55. Mock-infected mice were infused with saline using the same procedure (**Fig. 3A**). A higher HIV load (DNA or RNA copies) was detected in the plasma, spleen, and lateral hemisphere (LH) in ocln^−/−^ mice when compared with ocln^+/−^ or ocln^+/+^ mice (**Fig. 3B**). No changes in HIV load were found between genders (**Fig. 3C**).

**Figure 3.**
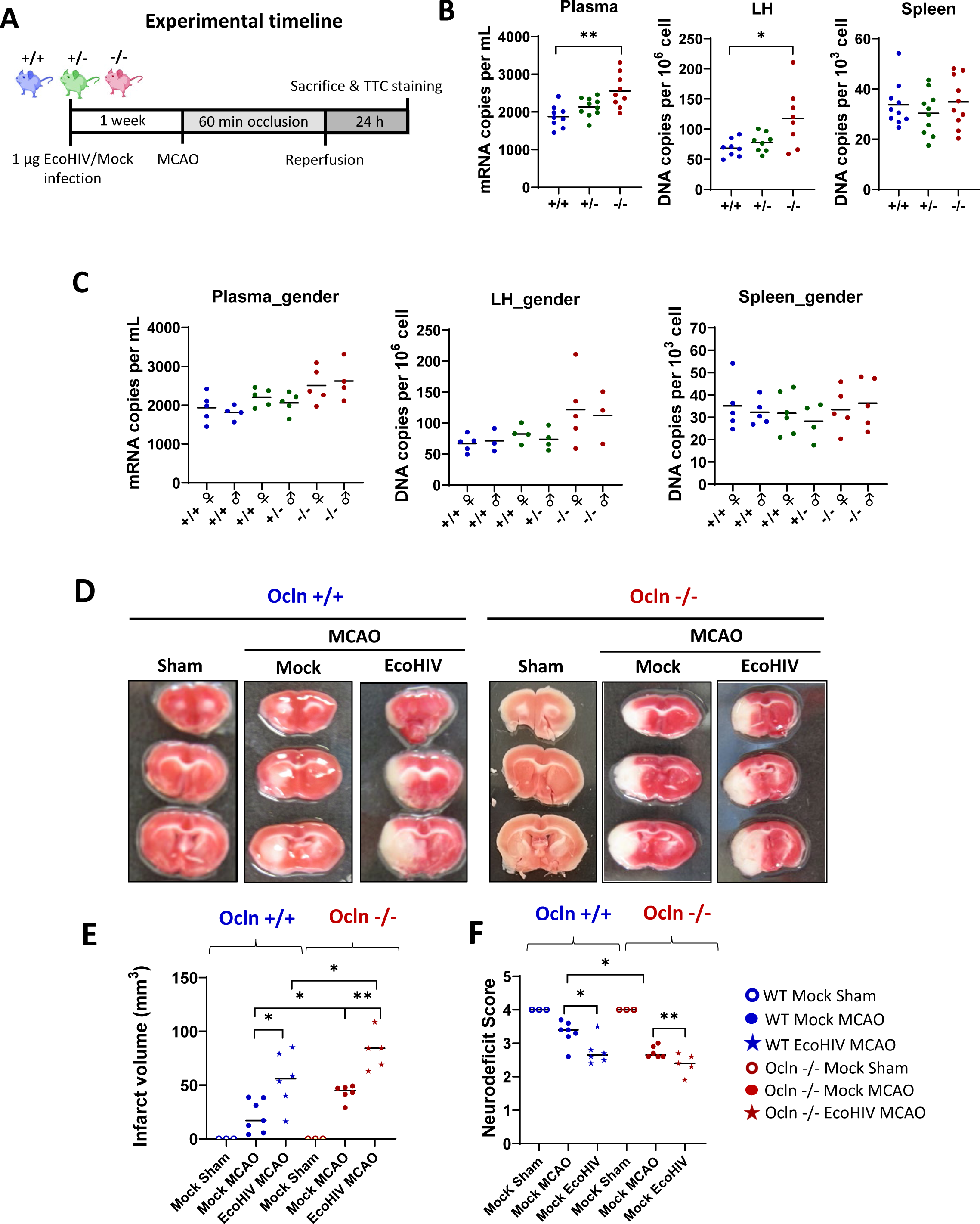
Ocln-deficient mice exhibit increased infarct size after EcoHIV infection. **(A)** A diagram reflecting the experimental timeline. Age- and sex-matched ocln^−/−^, ocln^+/−^, and ocln^+/+^ mice were inoculated with EcoHIV (1 µg of p24) via the internal carotid artery or Mock-infected. One week post-infection, ischemic stroke was induced by the MCAO for 24h, followed by reperfusion for 24h. **(B)** Levels of mRNA and DNA copies in plasma, lateral hemisphere (LH), and spleen were assessed by q-PCR. **(C)** mRNA and DNA copies from (B) as analyzed by gender of infected mice. **(D)** Representative staining of infarct lesions as determined by TTC staining of brain sections of ocln^−/−^ and ocln^+/+^ mice, mock, and EcoHIV-infected mice after ischemic stroke. Tissue damage is visualized as white areas. **(E)** Infarct size quantification from (D). **(F)** Assessment of neurological outcome 24h post-stroke. Individual data points are marked by blue (ocln^+/+^), green (ocln^+/−^), and red (ocln^−/−^) dots. \*\**p* = 0.003, \**p* < 0.0449, *n*= 3-11 animals per group, 7 independent experiments.

Knowing that ocln deficiency increases HIV infection, which is a risk factor for stroke, we then evaluated the outcomes of ischemic stroke in EcoHIV or Mock-infected ocln deficient mice. Briefly, ocln^−/−^, ocln^+/−^, and ocln^+/+^ mice were infected with EcoHIV, and stroke was induced by occlusion of the middle cerebral artery one week after infection (**Fig. 3A**). HIV infection resulted in a significant increase in stroke infarct volume as compared to mock-infected mice. Moreover, stroke volume was increased in ocln^−/−^ vs. ocln^+/−^, and ocln^+/+^ mice both in the mock and Eco-HIV groups (**Figs. 3D-E**). Next, we evaluated post-stroke motor functions by using a neurodeficit score 56, with the values between 0 (the lowest) and 4 (no motor deficit). Behavioral testing was assessed 24 h post-stoke. Eco-HIV-infected mice exhibited significant neurodeficits when compared with Mock-treated mice. In addition, ocln^−/−^ mice had significantly lower motor functionality when compared with ocln control mice (**Figs 3F**).

### Cellular ocln levels regulate the antiviral RIG-I pathway

Next, we focused on the mechanisms that may be responsible for ocln-mediated differential expression of the ISG genes and pathways by focusing on the expression of RIG-I, a critical pattern recognition receptor (PPR) in antiviral protection (**Fig. 4A**).

**Figure 4.**
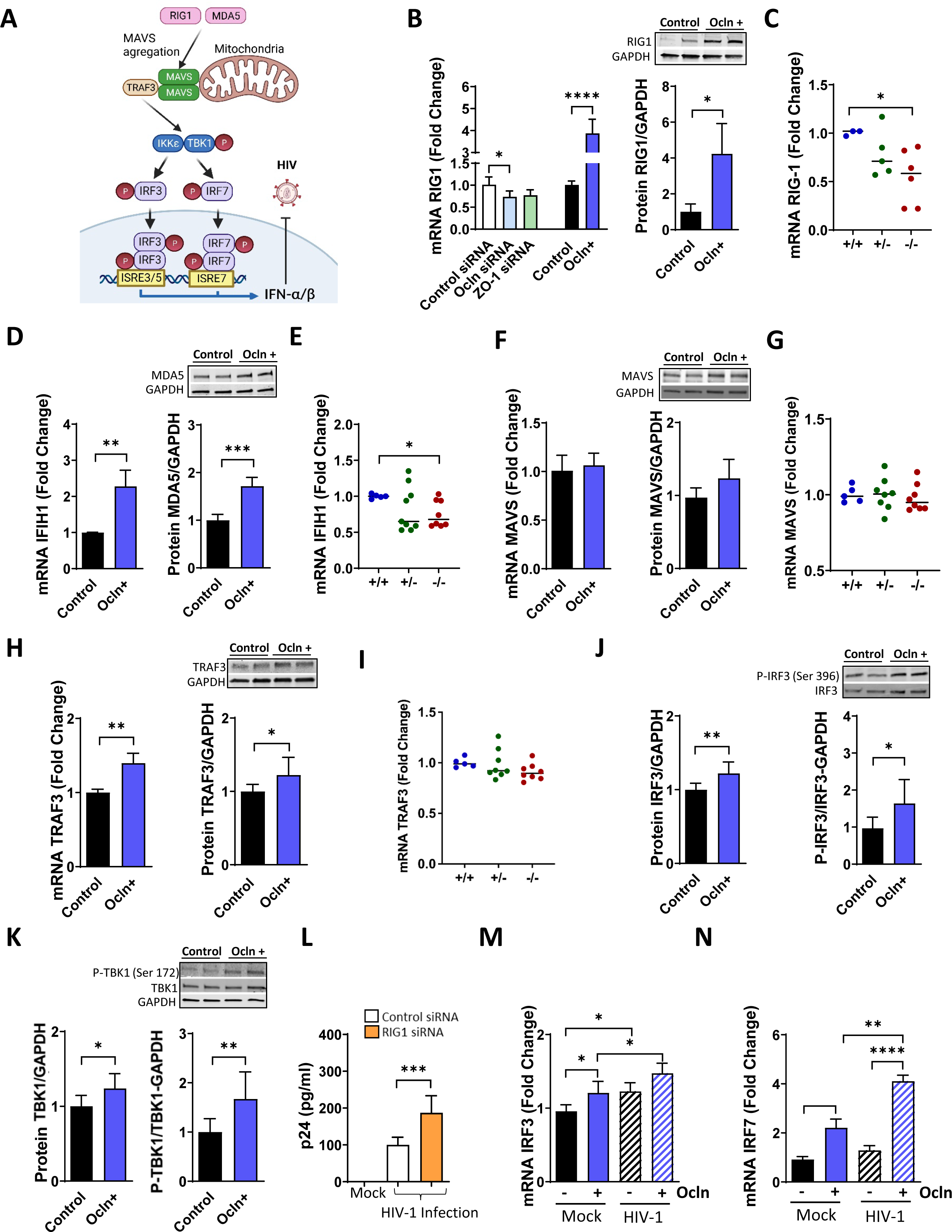
Ocln regulates the RIG-1-like receptor pathway signaling. **(A)** A diagram of antiviral RIG-1 pathway. Activated RIG-I and MDA5 molecules translocate to the mitochondria and interact with the MAVS adaptor. MAVS activates the downstream molecules TBK1/IKKε kinases, IRF3 and IRF7 inducing the transcriptional expression IFN-α/β. Pericytes were transfected with ocln-siRNA, ZO-1 siRNA, or PCMV3-OCLN vector as in Figure 2, and the expression of RIG-1 **(B)**, IFIH1 **(D)**, TRAF3 **(F)**, MAVS **(H)**, IRF3 and p-IRF3 **(J)**, and TBK1 and p-TBK1 **(K)** was evaluated by q-PCR and Western Blotting. **(L)** Pericytes were transfected with RIG-1-siRNA, and p24 levels were assessed as in Figure 2. **(M-N)** Pericytes were transfected with the PCMV3-OCLN vector and HIV-1 or mock-infected as in Figure 2. IRF3 and IRF7 mRNA levels were measured by q-PCR. n= 4-8 per group. **(C, E, G, I)** RNA was extracted from isolated microvessels of age- and sex-matched ocln^−/−^, ocln^+/−^, and ocln^+/+^ mice (n=5-11 animals per group) as in Figure 2. RIG-1 **(C)**, IFIH1 **(E)**, TRAF3 **(G)**, and MAVS **(I)** mRNA expression levels were measured by q-PCR. Graphs indicate the mean ± SD from three independent experiments. Individual animal data points are marked by blue (ocln^+/+^), green (ocln^+/−^), and red (ocln^−/−^) dots. ****p < 0.0001, ***p = 0.0002, **p = 0.003, *p < 0.0449.

The silencing of ocln markedly decreased the mRNA expression of RIG-I, with ocln overexpression having the opposite effects (**Fig. 4B**, left panel). Moreover, upregulation of cellular ocln levels significantly elevated RIG-I protein levels (**Fig. 4B**, right panel). While the percentage of changes in RIG-I expression was much higher after ocln overexpression compared to downregulation, these differences reflect a higher efficiency of ocln upregulation by the PCMV3-OCLN expression vector compared to downregulation by ocln silencing. We also evaluated RIG-I expression in cerebral microvessels isolated from the brains of ocln^+/+^ (controls), ocln^+/−^, and ocln^−/−^ mice. RIG-I mRNA levels were significantly lower in microvessels isolated from ocln-deficient mice as compared to controls with normal ocln expression (**Fig. 4C**). To confirm these results, we measured the input of cellular ocln levels on the expression of another PRR, namely MDA5, which is encoded by the IFH1 gene. Similar to changes in RIG-I, overexpression of ocln enhanced both IFH1 mRNA levels and MDA5 protein (**Fig. 4D)**. Moreover, IFH1 mRNA was decreased in microvessels of ocln-deficient mice (**Fig. 4E**). The importance of these findings stems from the fact that RIG-I and MDA5 are key molecules in the innate immune system. Specifically, they act as PRRs by recognizing pathogen-specific ligands, such as double stranded RNA and CpG DNA. PRRs can activate interferon regulatory factors (IRFs; IRF3 and 7) and induce the expression of type I IFNs 3,4,57. Downregulation of PRRs after ocln silencing is consistent with antiviral protection of this protein.

Because the signaling pathway downstream from RIG-I includes activation of MAVS and TNF receptor-associated factor (TRAF3) 3,5,7,58, we evaluated if ocln is involved in the regulation of these downstream proteins in BBB pericytes. Ocln overexpression did not affect MAVS expression levels; however, it significantly increased the expression of TRAF3 at mRNA and protein levels (**Figs. 4F and H**, respectively). No changes were found in TRAF3 and MAVS mRNA in microvessels of ocln-deficient mice when compared with control mice (**Figs. 4G, I).** We also investigated the downstream events of these events and analyzed changes in the phosphorylation of IRF3 and TBK1. Consistent with upregulation of TRAF3, both total protein levels and the levels of phosphorylated IRF3 (pIRF3) and phosphorylated TBK (pTBK) were elevated as the result of ocln overexpression **(Fig. 4J, K).**

In order to confirm the importance of these events in HIV infection, pericytes were transfected with RIG-I siRNA, followed by HIV infection. Diminished RIG-I expression markedly enhanced HIV replication **(Fig. 4L)** by ∼100% as measured by p24 levels in cell culture media and compared to cells transfected with control siRNA. Next, non-manipulated pericytes and pericytes with overexpressed ocln levels were either Mock-infected or infected with HIV-1 and the levels of IRF3 and 7 were analyzed. Upregulation of IRF3 mRNA levels in ocln-overexpressing cells was further increased when cells were infected with HIV-1 (**Fig. 4M**). Similar patterns of responses, but to a higher extent, were observed for IRF7 mRNA expression (**Fig. 4N)**. Altogether, these results indicate that ocln levels not only positively regulate the expression of RIG-I but also the downstream antiviral effectors of the RIG-I pathway.

### Ocln modulates mitochondrial dynamics and respiration

Knowing that a) mitochondrial dynamics is implicated in the regulation of the RIG-I signaling responses 14–17, b) MAVS is a mitochondria-associated protein and when activated, can influence mitochondrial dynamics and homeostasis 9,17, and c) ocln levels influences both the RIG-I signaling and MAVS activation (Fig. 4), we then evaluated if cellular ocln could influence the expression of the main mitochondrial dynamics regulatory proteins in human brain pericytes.

We first evaluated the impact of ocln on mitochondrial dynamics in the context of HIV infection by focusing on the genes that regulated mitochondria fusion and fission in human pericytes and cerebral microvessels of mice with altered ocln expression (**Fig. 5**). An increase in ocln levels resulted in a significant upregulation of mitochondrial fission regulatory protein FIS1 at both the mRNA and protein levels (**Fig. 5A**). Similarly, ocln overexpression increased MFF (mitochondria fission factor) mRNA (**Fig. 5C**) and mitochondrial fusion genes, such as MFN2 (mitofusin 2) and OPA1 (optic atrophy 1) (**Figs. 5E and 5G**, respectively). Except for an increase in FIS1 protein, no differences in protein levels of MFF, MFN2 and OPA1 **(Figs. 5C, E, G)** were found in pericytes overexpressing ocln. Additional HIV infection of pericytes did not result in altered expression of any of the studied genes related to mitochondrial fission and fusion in BBB pericytes. These results were confirmed in animal experiments. Microvessels isolated from the brains of mice deficient in ocln were characterized by decreased levels of FIS1 mRNA; however, no changes were found in MFF, MFN2 and OPA1 mRNA levels and infection with EcoHIV did not result in any changes in expression of this gene (**Figs. 5B, D, F, H**).

**Figure 5.**
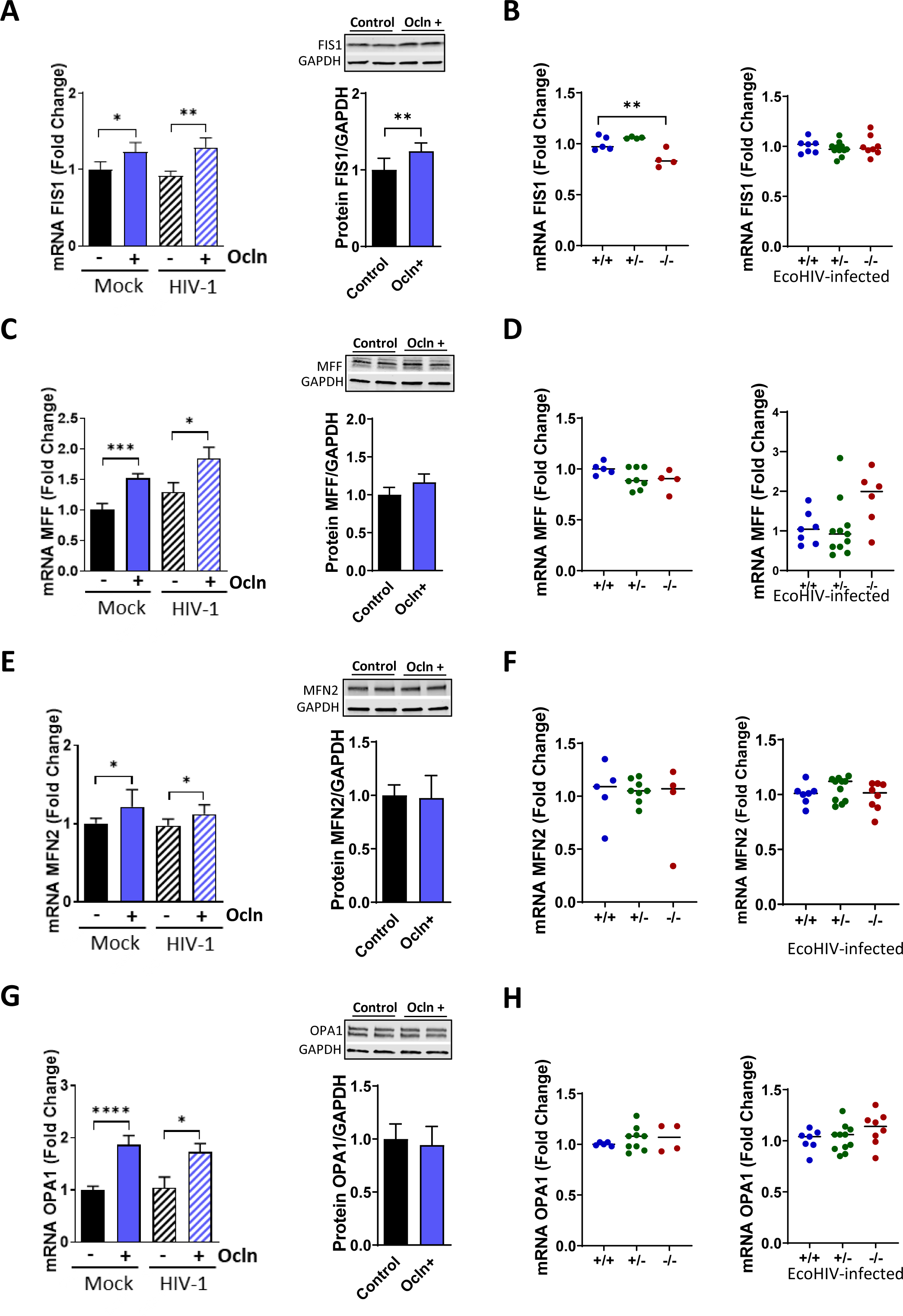
Impact of ocln on mitochondrial dysregulation. Pericytes were transfected with the PCMV3-OCLN vector and HIV-1 or mock-infected as in Figure 2. FIS1 **(A)**, MFF **(C)**, MFN2 **(E)**, and OPA1 **(G)** mRNA and protein levels were measured by q-PCR and Western Blotting. n= 4-8 per group. In addition, age- and sex-matched ocln^−/−^, ocln^+/−^, and ocln^+/+^ mice (n=4-11 animals per group) were EcoHIV or mock-infected and RNA was extracted from isolated microvessels. FIS1 **(B)**, MFF **(D)**, MFN2 **(F)** and OPA1 **(H)** mRNA expression levels were measured by q-PCR. Graphs indicate the mean ± SD from three independent experiments. Individual animal data points are marked by blue (ocln^+/+^), green (ocln^+/−^), and red (ocln^−/−^) dots. ****p < 0.0001, ***p = 0.0002, **p = 0.003, *p < 0.0449.

In the following series of experiments, we analyzed mitochondrial bioenergetics, metabolic stress, and subsequent ATP production in brain pericytes overexpressing different levels of ocln by using the Seahorse XF24 Analyzer and the Cell Mito Stress Test (**Figs. 6A-B**). Quantification of the oxygen consumption rate (OCR; **Fig. 6A**) indicated that there was an occludin-dependent decrease in basal respiration levels but no changes in non-mitochondria-related respiration when compared with control cells **(Fig. 6B)**. These results were consistent with a gradually lower maximal mitochondrial respiration, ATP-production, proton leak, and spare respiratory capacity in pericytes overexpressing different levels of ocln **(Fig. 6B)**. Overall, these novel results suggested that ocln overexpression suppresses mitochondrial oxidative electron transport chain-dependent respiration. Therefore, we then analyzed the impact of cellular ocln levels on mitochondrial morphology by performing TOM20 immunostaining in pericytes overexpressing ocln (**Fig. 6C**). Upregulation of ocln resulted in a significant decrease in the mitochondrial branch length (**Fig. 6D**), with no changes in mitochondrial footprint analysis (**Fig. 6E**). Nevertheless, an increase in ocln levels resulted in a significant elevation of mitochondrial ROS as assessed by MitoSox Red (**Fig. 6F**). No changes were found in the overall protein levels of TOM20, a general marker of mitochondrial mass (**Fig. 6G**).

**Figure 6.**
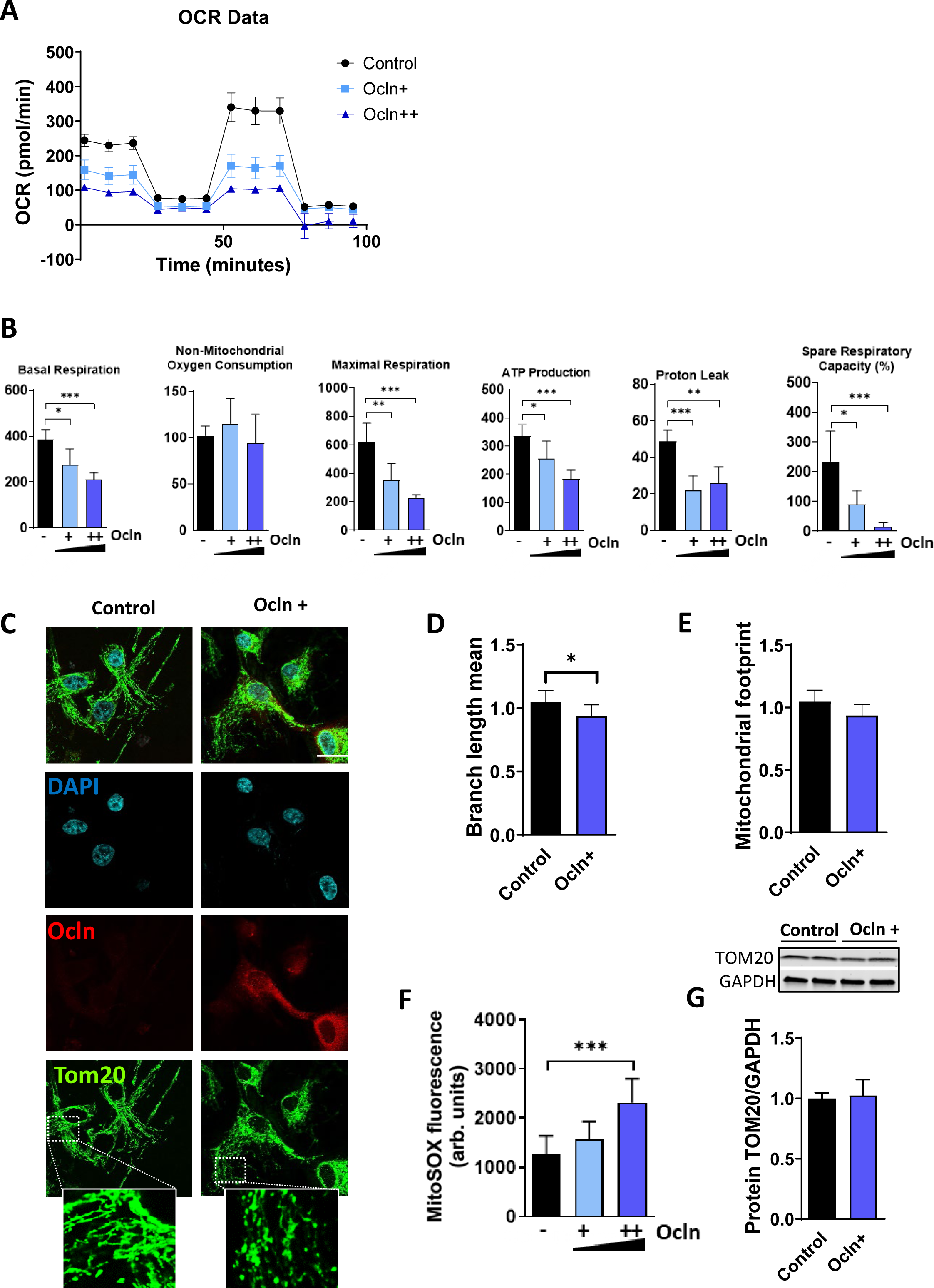
Impact of ocln on mitochondrial respiratory function. **(A)** Seahorse Cell Mito Stress Test analysis of OCR after transfecting pericytes with different concentrations of PCMV3-OCLN vector. **(B)** Quantitative measurements of basal respiration, non-mitochondrial oxygen consumption, maximal respiration, ATP production, proton leak, and spare respiration capacity were performed. **(C)** Confocal microscopy images of pericytes after ocln overexpression. Cells were stained with DAPI (blue), ocln (red.), and TOM20 (green.) for tracking the mitochondrial membrane. **(D-E)** Mitochondria was skeletonized, and mitochondrial mean branch **(D)** and footprint **(E)** were quantified by using the MiNA plug-in on Image J**. (F)** Measurement of mitochondrial superoxide as determined by MitoSOX Red after transfecting pericytes with different concentrations of PCMV3-OCLN vector. Fluorescence intensity of MitoSOX Red was normalized to protein levels. **(G)** Total TOM20 protein levels were measured by Western Blotting. Graphs indicate the mean ± SD from three independent experiments. ***p = 0.0002, **p = 0.003, *p < 0.0449, n= 4-8 per group; scale bars, 30 μm. OCR: oxygen consumption rate; MiNA: mitochondrial network analysis; DAPI: 4′,6-diamidino-2-phenylindole.

### Ocln promotes cellular autophagy and apoptosis

Dysregulation in mitochondrial function has been linked to alterations in autophagy and apoptosis 59–62 pathways. Additionally, an association between activation of the RIG-I signaling pathway, MAVS, and autophagy has been reported 12,13,63. Therefore, we investigated whether the modulation of ocln expression can alter autophagy and/or apoptosis in human brain pericytes. Autophagy activation was assessed by analyzing the protein levels of microtubule-associated protein 1 light chain 3B (LC3B) and the conversion of LC3B-I to LC3B-II, p62/sequestosome 1 (SQSTM1), and the autophagic receptor NDP52.

After autophagy induction, the cytoplasmic form of LC3 (LC3-I) is cleaved and translocates to the membrane, forming LC3-II and joining with autophagic vesicles 64–66. When compared to controls, LC3-I conversion to LC3-II was significantly enhanced after ocln upregulation, indicating autophagosome formation (**Fig. 7A**). In line with these results, p62, a protein known to interact with L3CB and to promote ubiquitination of target substrates in autophagosomes 67,68, was also increased at both mRNA and protein levels in ocln overexpressing cells (**Fig. 7B**). Consistently, the expression of the gene CALCOCO2, which encodes for the Nuclear Domain 10 Protein 52 (NLP52), a receptor implicated in directing autophagy targets to autophagosomes 69, was increased at the mRNA and protein levels (**Fig. 7C**). The peroxisome proliferator-activated receptor gamma coactivator 1-alpha (PGC-1α), a transcriptional coactivator involved in mitochondrial biogenesis 70,71, was upregulated at the protein levels in ocln-overexpressing pericytes. In contrast, the expression of the gene PPARGC1A, which encodes for PGC-1α was unchanged in these cells at the studied time point (**Fig. 7D**).

**Figure 7.**
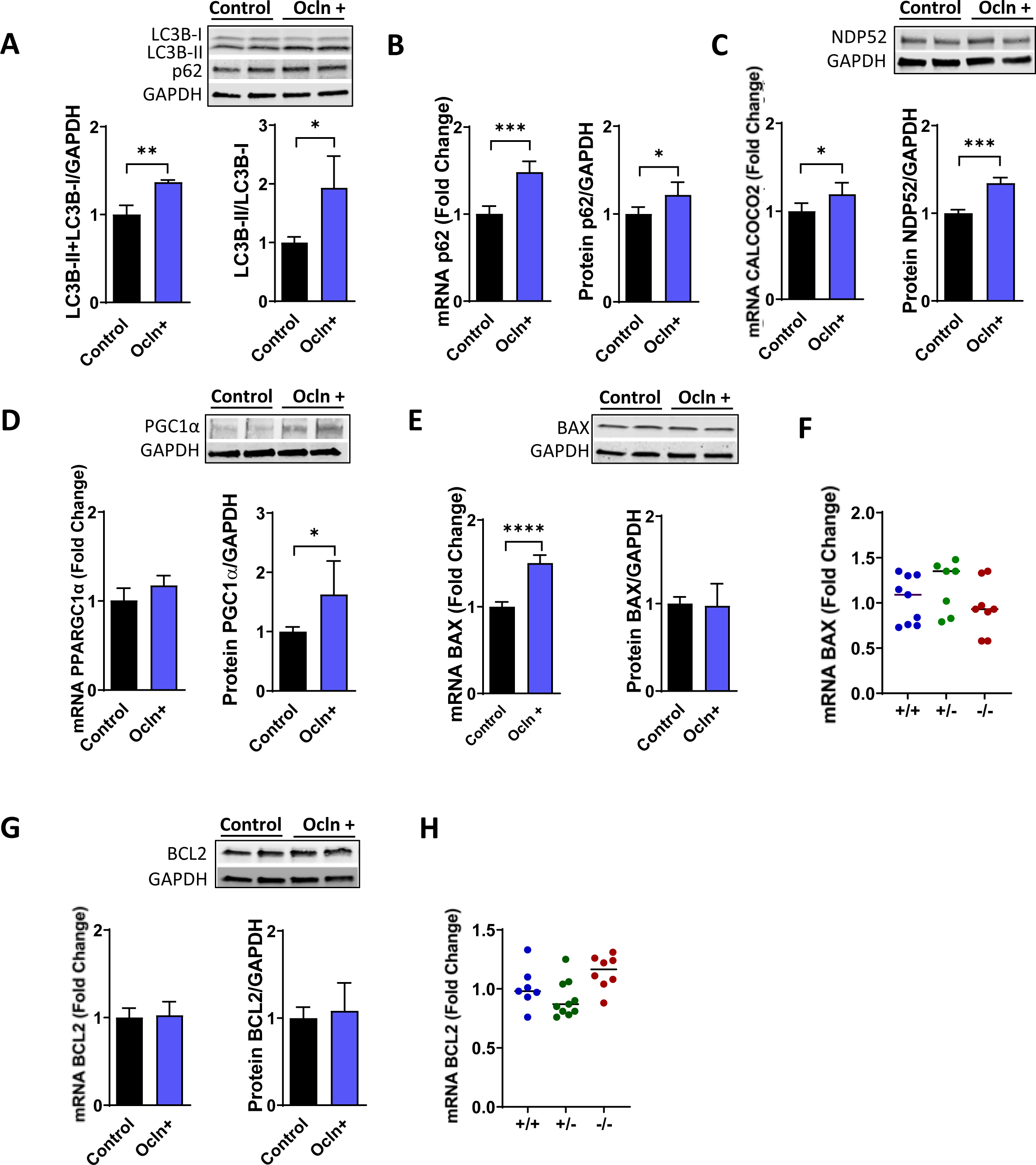
Impact of ocln on autophagy and apoptosis regulation. Pericytes were transfected with the PCMV3-OCLN vector as in Figure 2. LC3B-I and LC3B-II **(A)**, p62 **(B)**, NDP52 **(C)**, PGC1α **(D)**, BAX **(E)**, and BCL2 **(G)** mRNA and protein levels were measured by q-PCR and Western Blotting. n=4-8 per group. BAX **(F)** and BCL2 **(H)** mRNA expression levels from extracted microvessels isolated from age- and sex-matched ocln^−/−^, ocln^+/−^, and ocln^+/+^ mice (n=7-10 animals per group). Graphs indicate the mean ± SD from three independent experiments. Individual animal data points are marked by blue (ocln^+/+^), green (ocln^+/−^), and red (ocln^−/−^) dots. ****p < 0.0001, ***p = 0.0002, **p = 0.003, *p < 0.0449.

The expression levels of proapoptotic BAX and antiapoptotic BCL2 genes and proteins were assessed by q-PCR and immunoblotting as early markers of apoptosis. Even though ocln overexpression increased the BAX mRNA expression, this effect was not observed at the protein levels (**Fig. 7E**). Moreover, BAX protein levels were unchanged in microvessels isolated from the brains of ocln-deficient mice as compared to controls (**Fig. 7F**). No differences were found at the mRNA and protein levels of BCL2 in ocln-overexpressing pericytes or ocln-deficient mice (**Fig. 7G-H**). Taken together, these results suggest that ocln can alter mitochondrial dynamics and modulate autophagy, with a limited impact on pro-apoptotic-associated responses.

## DISCUSSION

Emerging evidence indicates that occludin is a multifunctional protein with a key role in regulating human brain pericyte metabolism. Our laboratory has previously shown that ocln can act as a NADH oxidase and modulate AMPK protein kinase activity in BBB pericytes 31,43. Taking the importance of cellular ocln levels into consideration, we performed a transcriptome analysis to identify changes in gene expression related to ocln downregulation. Our results **(Fig. 1)** identified that among differentially altered cellular pathways, eleven were related to IFN signaling. The importance of these observations stem from the fact that IFN signaling and ISG expression levels have been associated with antiviral functions in several viral infections, including HIV-1 21-24,72.

While the majority of HIV-1 replication in the brain appears to occur in microglial cells and perivascular macrophages 73–75, growing evidence indicates that specific cell types of the BBB, such as astrocytes 74,76 and brain pericytes 30–35 can be infected by HIV-1. Canonical HIV-1 infection uses both the main HIV-1 receptor, CD4, and the co-receptors, primarily CXCR4 and CCR5. Therefore, it is important that pericytes express high levels of HIV-1 co-receptors, CCR5 and CXCR4, and also CD4 77. Importantly, HIV-1-mediated alterations of ocln expression have been linked to the regulatory mechanisms of HIV-1 infection 25,31,35. Specifically, a decrease in cellular ocln levels results in enhanced HIV-1 replication, and the opposite effect is observed upon overexpression of ocln 31,35. At least part of this effect is related to ocln having NADH oxidase activity, which can influence the expression and activation of the class-III histone deacetylase SIRT-1 31. Moreover, ocln can regulate HIV-1 budding from the infected cells by forming a complex with caveolin-1 and ALIX 35. While the majority of research on the regulatory role of ocln on HIV-1 infection was performed on brain pericytes 25,31,35, the findings were confirmed in human primary macrophages, differentiated monocytic U937 cells, and HEK-293 cells 43.

The findings of the present study indicate that ocln levels can regulate HIV-1 infection by impairing the function of ISGs 78,79. Indeed, we present novel results that ocln levels can regulate the expression of several ISGs and/or proteins such as ISG15, MX1, MX2, IFIT1, HERC5, and USP18. In support of the findings on the interrelationship between ocln and ISG, we demonstrated that ocln can regulating the OAS ISG family in BBB pericytes 25. Moreover, a study has been published showing that IL-22 could increase MX2 and ocln expression levels in end1/E6E7cells 80. Confirming the impact of ocln on the IFN-related pathways, our novel results indicated that the expression levels of IFNα5, IFNβ1, IFNɣ, and ISG15 were significantly lower in ocln^−/−^ mice when compared with ocln^+/+^ mice (**Fig. 2**).

Our novel findings demonstrated that downregulation of ISG15 and MX2 can result in a significant increase in the rate of HIV-1 infection in pericytes, as determined by p24 levels. Supporting these findings, it was shown that lower levels of ISG15 enhance HIV-1 replication 22, and MX2 has been recognized as a key host factor protecting against HIV-1 replication 21,81,82. Conversely, despite potent antiviral activity of MX1 against several viruses, such as the Thogoto virus 83, hepatitis B virus 84, and influenza A virus 85, its role in HIV-1 infection remains controversial, with several studies failing to show any HIV-1-specific antiviral activity 86. In line with these reports, our results indicate that silencing MX1 did not affect HIV-1 replication in pericytes (**Fig. 2**).

The impact of ocln on HIV infection was next confirmed in animal studies in ocln-deficient mice. Mice were infected with EcoHIV for one week, followed by the measurement of p24 levels in plasma, spleen, and LH. We indicated for the first time that ocln^−/−^ mice expressed a significantly higher number of copies of mRNA and DNA compared to ocln^+/−^ or ocln^+/+^ mice. However, no changes in HIV load were found when comparing between genders (**Fig. 3**). Several studies have described that the risk and the severity of cerebrovascular diseases, and particularly ischemic stroke, are significantly higher in people living with HIV 87. Since ocln deficiency increases HIV infection, we next evaluated the outcomes of ischemic stroke in EcoHIV or Mock-infected ocln-deficient mice. The present study is the first to demonstrate that ocln^−/−^ mice display a significantly larger infract volume after HIV infection when compared to ocln^+/+^-infected mice (**Fig. 3**). These results are important since little is known about the mechanisms by which HIV worsens cerebrovascular diseases such as ischemic stroke.

The present study also reports that the RIG-I and MDA5 genes, two key molecules in the innate immune system, were altered as the result of ocln overexpression in human brain pericytes. Moreover, our data show that ocln can regulate RIG-1 and MDA5 *in vivo*. RIG-I and MDA5 mRNA levels were significantly lower in microvessels isolated in ocln-deficient mice as compared to controls with normal ocln expression. Both RIG-I and MDA5 are cytosolic pattern recognition receptors (PRR) involved in the IFN1-mediated responses after virus infection 3,4,7,58. Following the downstream pathway, ocln overexpression also activated MAVS and induced TBK1 and IRF3 phosphorylation. Interestingly, there was also upregulation of IRF3 and IRF7 in response to ocln overexpression (**Fig. 4**), with this effect being increased when cells were infected with HIV-1. These results are consistent with the notion that IRF3 and IRF7 are key regulators of type I IFN gene expression in response to viral infections 88. Although IRF3 has limited DNA binding, it can induce the transcription of IFNβ, IFNα4 and IFNα1 genes. Moreover, IRF3 can bind to IRF7, regulating in concert IFNα/β induction. IRF7 has broader DNA specificity and is able to induce the production of several IFNs, including IFNα2, IFNα5, IFNα6, INFα8, IFNβ2 and IFNβ3 89,90. These results are novel since no previous studies have shown a relationship between ocln and the antiviral RIG-I signaling pathway.

Knowing that MAVS activation can influence mitochondrial dynamics 9,17, while ocln can alter RIG-1 like receptor signaling, we then evaluated the role of ocln in mitochondrial dynamics, a process that allows mitochondria to regulate and maintain their integrity and homeostasis. Mitochondrial dynamics is established through the processes of fusion and fission where dysregulation of this process has been linked to several diseases such as neurodegenerative and cardiovascular diseases 91. We indicated that modulation of ocln resulted in significant changes in the expression of FIS1 at mRNA and protein levels both *in vitro* and *in vivo* (**Fig. 5**), a key protein in maintaining mitochondrial fission 92. This dysregulation of mitochondrial dynamics was associated with impaired mitochondrial bioenergetics (**Fig. 6**), indicating that alterations of ocln levels can induce mitochondrial stress and metabolic dysfunction. This conclusion is in line with the observations that ocln can promote AMPK expression and activation, influencing the expression of glucose transporters GLUT-1 and GLUT-4, glucose uptake, and ATP content 43.

Dysregulation in mitochondrial dynamics has been linked to autophagy and apoptosis 59–62. While ocln overexpression elevated the expression of cellular markers of autophagy, it had only limited impact on activation of apoptosis, suggesting dysregulation of these processes (**Figs. 7**). Although some literature reports have linked ocln levels with both autophagy activation 93–95 and apoptosis 96–100, the results are controversial, and this relationship was never studied in microvasculature. In contrast to our findings, a reduction in susceptibility to the induction of apoptosis in keratinocytes after ocln downregulation has been previously reported 96.

In conclusion, the present study reveals the significant impact of cellular ocln levels on the modulation of autoimmune responses and anti-HIV signaling through the regulation of ISGs, RIG-I, and MAVS activation. Cellular ocln levels also influence mitochondrial dynamics and bioenergetics, as well as autophagy. Overall, these findings establish ocln as an important regulator of innate immune responses in the context of cerebrovascular pathology associated with HIV infection, including ischemic stroke.

## Acknowledgments

Supported by the National Institutes of Health grants HL126559, MH128022, MH122235, MH072567, DA050528, DA044579, and DA059849. ST was in part supported by the National Institute of Allergy and Infectious Diseases of the National Institutes of Health (NIAID), grant number P30AI073961 and by an AHA postdoctoral fellowship (20POST35211181). Olivia M. Osborne was in part supported by the National Institute of Neurological Disorders and Stroke (NINDS) and the National Institute of Aging (NIA) of the National Institutes of Health under award number NS125905.

## Author contributions

S.T. and M.T. designed the study. S.T., S.R., O.N., T.T., and O.O., performed and analyzed the data from gene silencing and overexpression, western blots, RT-qPCR, immunofluorescence staining and mitochondrial potential/oxidative stress measurements. S.T. and O.N., produced the viral stocks and performed all ELISA assays. S.T., M. P., and E.S performed EcoHIV infection in mice and MCAO surgery. T.M performed all RNA sequencing data analysis. S.T drafted the manuscript. M.T. edited and revised the manuscript. All co-authors approved the final version of the manuscript.

## Declaration of Interests

The authors declare no competing interests.

## Inclusion and Diversity

One or more of the authors of this paper self-identifies as an underrepresented ethnic minority in science. One or more of the authors of this paper received support from a program designed to increase minority representation in science.

## Data availability

All source data supporting the findings of this manuscript are available from the corresponding authors upon request.

---

**Supplemental Table 1.** Genes affected in each Reactome pathway after ocln silencing in human brain pericytes.

| Pathway | Genes |
| --- | --- |
| **Interferon alpha/beta signaling** | IFI35, XAF1, HLA-F, , ISG20, IRF7, BST2, IFI6, IFI27, SAMHD1, USP18, IFITM1, ISG15, OAS3,IFIT3,IFIT1, OAS2, OAS1, IFIT2, MX1, MX2, OASL, RSAD2 |
| **Interferon Signaling** | GBP4, STAT1, IFI35, XAF1, HLA-F, ISG20, IRF7, BST2, IFI6, IFI27, SAMHD1, DDX58, USP18 , IFITM1, ISG15, OAS3, IFIT3, IFIT1, OAS2, OAS1, IFIT2, HERC5, MX1, MX2, OASL, RSAD2 |
| **Cytokine Signaling in Immune system** | KIT, GBP4, STAT1, IFI35, CCL5, XAF1, TNFSF13B, LGALS9, HLA-F, ISG20, IRF7, BST2, IFI6, IFI27, SAMHD1, DDX58, USP18, IFITM1, ISG15, OAS3, IFIT3, IFIT1, OAS2, CXCL10, OAS1, IFIT2, HERC5, MX1, MX2, OASL, RSAD2 |
| **Antiviral mechanism by IFN-stimulated genes** | STAT1, DDX58, USP18, ISG15, OAS3, IFIT1, OAS2, OAS1, HERC5, MX1, MX2,OASL |
| **OAS antiviral response** | DDX58, OAS3, OAS2, OAS1, OASL |
| **Immune System** | KIT, DTX3L, GBP4, STAT1, IFI35, CCL5, DHX58, XAF1, TNFSF13B, LGALS9, HLA-F, ISG20, IRF7, BST2, IFI6, IFI27, SAMHD1, DDX58, USP18, IFITM1, IFIH1, HERC6, ISG15, OAS3, IFIT3, IFIT1, OAS2, CXCL10, OAS1, IFIT2, HERC5, MX1, MX2, OASL, RSAD2 |
| **ISG15 antiviral mechanism** | STAT1, DDX58, USP18, ISG15, IFIT1, HERC5, MX1, MX2 |
| **Interferon gamma signaling** | GBP4, HLA-F, IRF7, OAS3, OAS2, OAS1, OASL |
| **DDX58/IFIH1-mediated induction of interferon-alpha/beta** | DHX58, IRF7, DDX58, IFIH1, ISG15, HERC5 |
| **Negative regulators of DDX58/IFIH1 signaling** | DDX58, IFIH1, ISG15, HERC5 |
| **TRAF3-dependent IRF activation pathway** | IRF7, DDX58, IFIH1 |

**Supplemental Figure 1.**
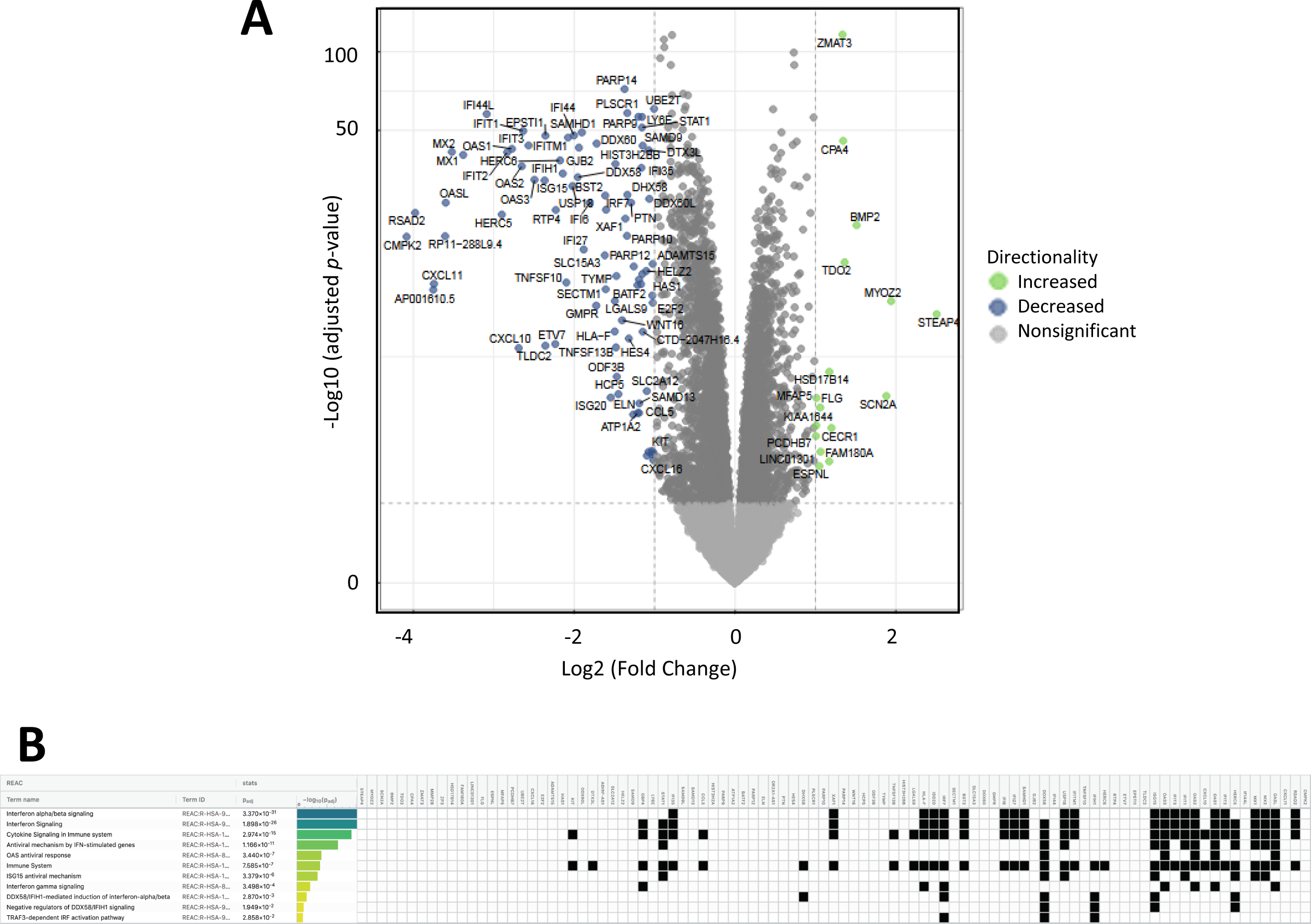
Ocln silencing displays distinct RNA-seq transcriptome signatures. Related to Figure 1. **(A)** Volcano plot highlighting the differentially expressed genes in pericytes with silenced ocln and controls. Blue dots represent downregulated genes; green dots, upregulated genes, and grey dots genes that were not significantly affected. **(B)** Gene set enrichment analysis (GSEA) was performed using the g: Profiler and Reactome pathways. Figure show the *p*-values for the Reactome-enriched pathways analysis.

**Supplemental Figure 2.**
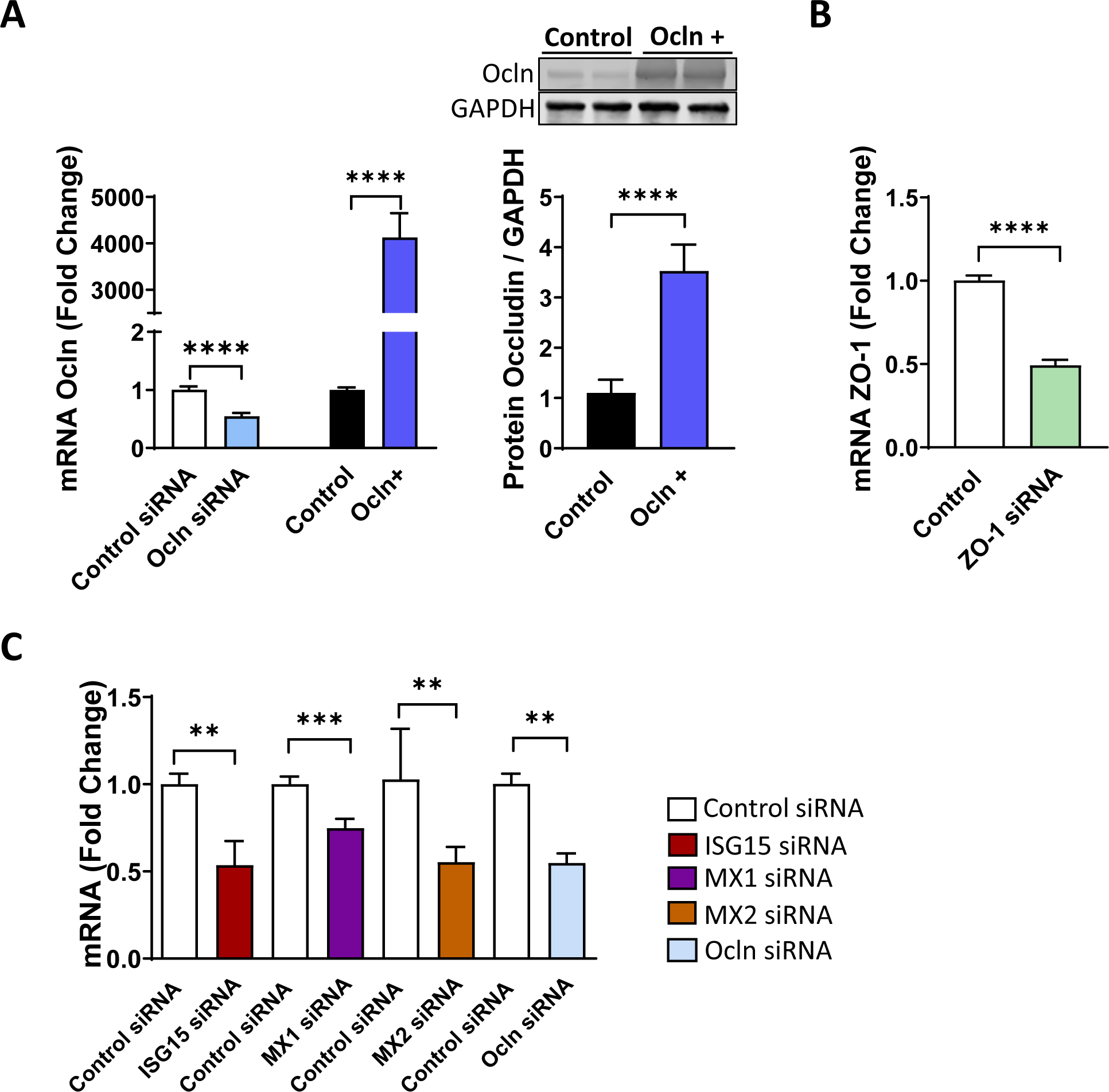
Effectiveness of silencing procedures employed in the study. Related to Figure 2. Pericytes were transfected with ocln-siRNA, ZO-1 siRNA, or PCMV3-OCLN vectors, and the expression of genes encoding for ocln **(A)** and ZO-1 **(B)** was evaluated by q-PCR and Western Blotting. **(C)** Pericytes were transfected with ISG15 siRNA, MX1 siRNA, MX2 siRNA, and ocln siRNA. The mRNA expression levels of ISG15, MX1, MX2, and ocln were evaluated by q-PCR. Graphs indicate the mean ± SD from three independent experiments. ****p < 0.0001, ***p = 0.0002, **p = 0.003, n=3-8 per group.

## Notes

### Competing Interest Statement

The authors have declared no competing interest.

## References

1. Belgnaoui, S.M., Paz, S., and Hiscott, J. (2011). Orchestrating the interferon antiviral response through the mitochondrial antiviral signaling (MAVS) adapter. Curr Opin Immunol 23, 564–572. 10.1016/j.coi.2011.08.001.

2. Yoneyama, M., Kikuchi, M., Natsukawa, T., Shinobu, N., Imaizumi, T., Miyagishi, M., Taira, K., Akira, S., and Fujita, T. (2004). The RNA helicase RIG-I has an essential function in double-stranded RNA-induced innate antiviral responses. Nat Immunol 5, 730–737. 10.1038/ni1087.

3. Rice, M., Tili, E., Loghmani, H., and Nuovo, G.J. (2023). The differential expression of toll like receptors and RIG-1 correlates to the severity of infectious diseases. Annals of diagnostic pathology 63, 152102. 10.1016/j.anndiagpath.2022.152102.

4. Vlaming, K.E., van Wijnbergen, K., Kaptein, T.M., Nijhuis, M., Kootstra, N.J., de Bree, G.J., and Geijtenbeek, T.B. (2023). Crosstalk between TLR8 and RIG-I-like receptors enhances antiviral immune responses. Frontiers in medicine 10, 1146457. 10.3389/fmed.2023.1146457.

5. Hou, F., Sun, L., Zheng, H., Skaug, B., Jiang, Q.X., and Chen, Z.J. (2011). MAVS forms functional prion-like aggregates to activate and propagate antiviral innate immune response. Cell 146, 448–461. 10.1016/j.cell.2011.06.041.

6. Qi, N., Shi, Y., Zhang, R., Zhu, W., Yuan, B., Li, X., Wang, C., Zhang, X., and Hou, F. (2017). Multiple truncated isoforms of MAVS prevent its spontaneous aggregation in antiviral innate immune signalling. Nat Commun 8, 15676. 10.1038/ncomms15676.

7. Zheng, Y., Zhuang, M.W., Han, L., Zhang, J., Nan, M.L., Zhan, P., Kang, D., Liu, X., Gao, C., and Wang, P.H. (2020). Severe acute respiratory syndrome coronavirus 2 (SARS-CoV-2) membrane (M) protein inhibits type I and III interferon production by targeting RIG-I/MDA-5 signaling. Signal transduction and targeted therapy 5, 299. 10.1038/s41392-020-00438-7.

8. Tu, D., Zhu, Z., Zhou, A.Y., Yun, C.H., Lee, K.E., Toms, A.V., Li, Y., Dunn, G.P., Chan, E., Thai, T., et al. (2013). Structure and ubiquitination-dependent activation of TANK-binding kinase 1. Cell Rep 3, 747–758. 10.1016/j.celrep.2013.01.033.

9. Seth, R.B., Sun, L., Ea, C.K., and Chen, Z.J. (2005). Identification and characterization of MAVS, a mitochondrial antiviral signaling protein that activates NF-kappaB and IRF 3. Cell 122, 669–682. 10.1016/j.cell.2005.08.012.

10. Tal, M.C., and Iwasaki, A. (2011). Mitoxosome: a mitochondrial platform for cross-talk between cellular stress and antiviral signaling. Immunol Rev 243, 215–234. 10.1111/j.1600-065X.2011.01038.x.

11. Sun, Q., Sun, L., Liu, H.H., Chen, X., Seth, R.B., Forman, J., and Chen, Z.J. (2006). The specific and essential role of MAVS in antiviral innate immune responses. Immunity 24, 633–642. 10.1016/j.immuni.2006.04.004.

12. Lin, Y., Huang, C., Gao, H., Li, X., Lin, Q., Zhou, S., Huo, Z., Huang, Y., Liu, C., and Zhang, P. (2022). AMBRA1 promotes dsRNA- and virus-induced apoptosis through interacting with and stabilizing MAVS. Journal of cell science 135. 10.1242/jcs.258910.

13. Liang, R., Song, H., Wang, K., Ding, F., Xuan, D., Miao, J., Fei, R., and Zhang, J. (2022). Porcine epidemic diarrhea virus 3CL(pro) causes apoptosis and collapse of mitochondrial membrane potential requiring its protease activity and signaling through MAVS. Veterinary microbiology 275, 109596. 10.1016/j.vetmic.2022.109596.

14. Yasukawa, K., Oshiumi, H., Takeda, M., Ishihara, N., Yanagi, Y., Seya, T., Kawabata, S., and Koshiba, T. (2009). Mitofusin 2 inhibits mitochondrial antiviral signaling. Science signaling 2, ra47. 10.1126/scisignal.2000287.

15. Castanier, C., Garcin, D., Vazquez, A., and Arnoult, D. (2010). Mitochondrial dynamics regulate the RIG-I-like receptor antiviral pathway. EMBO reports 11, 133–138. 10.1038/embor.2009.258.

16. Berry, N., Suspene, R., Caval, V., Khalfi, P., Beauclair, G., Rigaud, S., Blanc, H., Vignuzzi, M., Wain-Hobson, S., and Vartanian, J.P. (2021). Herpes Simplex Virus Type 1 Infection Disturbs the Mitochondrial Network, Leading to Type I Interferon Production through the RNA Polymerase III/RIG-I Pathway. mBio 12, e0255721. 10.1128/mBio.02557-21.

17. Li, S., Kuang, M., Chen, L., Li, Y., Liu, S., Du, H., Cao, L., and You, F. (2021). The mitochondrial protein ERAL1 suppresses RNA virus infection by facilitating RIG-I-like receptor signaling. Cell reports 34, 108631. 10.1016/j.celrep.2020.108631.

18. Fenton-May, A.E., Dibben, O., Emmerich, T., Ding, H., Pfafferott, K., Aasa-Chapman, M.M., Pellegrino, P., Williams, I., Cohen, M.S., Gao, F., et al. (2013). Relative resistance of HIV-1 founder viruses to control by interferon-alpha. Retrovirology 10, 146. 10.1186/1742-4690-10-146.

19. Marsili, G., Remoli, A.L., Sgarbanti, M., Perrotti, E., Fragale, A., and Battistini, A. (2012). HIV-1, interferon and the interferon regulatory factor system: an interplay between induction, antiviral responses and viral evasion. Cytokine Growth Factor Rev 23, 255–270. 10.1016/j.cytogfr.2012.06.001.

20. Nazli, A., Dizzell, S., Zahoor, M.A., Ferreira, V.H., Kafka, J., Woods, M.W., Ouellet, M., Ashkar, A.A., Tremblay, M.J., Bowdish, D.M., and Kaushic, C. (2019). Interferon-beta induced in female genital epithelium by HIV-1 glycoprotein 120 via Toll-like-receptor 2 pathway acts to protect the mucosal barrier. Cell Mol Immunol 16, 178–194. 10.1038/cmi.2017.168.

21. Betancor, G. (2023). You Shall Not Pass: MX2 Proteins Are Versatile Viral Inhibitors. Vaccines 11. 10.3390/vaccines11050930.

22. Osei Kuffour, E., Konig, R., Haussinger, D., Schulz, W.A., and Munk, C. (2019). ISG15 Deficiency Enhances HIV-1 Infection by Accumulating Misfolded p53. mBio 10. 10.1128/mBio.01342-19.

23. Michalska, A., Blaszczyk, K., Wesoly, J., and Bluyssen, H.A.R. (2018). A Positive Feedback Amplifier Circuit That Regulates Interferon (IFN)-Stimulated Gene Expression and Controls Type I and Type II IFN Responses. Frontiers in immunology 9, 1135. 10.3389/fimmu.2018.01135.

24. Yi, D.R., An, N., Liu, Z.L., Xu, F.W., Raniga, K., Li, Q.J., Zhou, R., Wang, J., Zhang, Y.X., Zhou, J.M., et al. (2019). Human MxB Inhibits the Replication of Hepatitis C Virus. Journal of virology 93. 10.1128/JVI.01285-18.

25. Torices, S., Teglas, T., Naranjo, O., Fattakhov, N., Frydlova, K., Cabrera, R., Osborne, O.M., Sun, E., Kluttz, A., and Toborek, M. (2023). Occludin Regulates HIV-1 Infection by Modulation of the Interferon Stimulated OAS Gene Family. Molecular neurobiology, 1-17. 10.1007/s12035-023-03381-0.

26. Bell, R.D., Winkler, E.A., Sagare, A.P., Singh, I., LaRue, B., Deane, R., and Zlokovic, B.V. (2010). Pericytes control key neurovascular functions and neuronal phenotype in the adult brain and during brain aging. Neuron 68, 409–427. 10.1016/j.neuron.2010.09.043.

27. Kamouchi, M., Ago, T., and Kitazono, T. (2011). Brain pericytes: emerging concepts and functional roles in brain homeostasis. Cell Mol Neurobiol 31, 175–193. 10.1007/s10571-010-9605-x.

28. Iwao, T., Takata, F., Matsumoto, J., Goto, Y., Aridome, H., Yasunaga, M., Yokoya, M., Kataoka, Y., and Dohgu, S. (2023). Senescence in brain pericytes attenuates blood-brain barrier function in vitro: A comparison of serially passaged and isolated pericytes from aged rat brains. Biochemical and biophysical research communications 645, 154–163. 10.1016/j.bbrc.2023.01.037.

29. Gurler, G., Soylu, K.O., and Yemisci, M. (2022). Importance of Pericytes in the Pathophysiology of Cerebral Ischemia. Noro psikiyatri arsivi 59, S29–S35. 10.29399/npa.28171.

30. Nakagawa, S., Castro, V., and Toborek, M. (2012). Infection of human pericytes by HIV-1 disrupts the integrity of the blood-brain barrier. J Cell Mol Med 16, 2950–2957. 10.1111/j.1582-4934.2012.01622.x.

31. Castro, V., Bertrand, L., Luethen, M., Dabrowski, S., Lombardi, J., Morgan, L., Sharova, N., Stevenson, M., Blasig, I.E., and Toborek, M. (2016). Occludin controls HIV transcription in brain pericytes via regulation of SIRT-1 activation. FASEB J 30, 1234–1246. 10.1096/fj.15-277673.

32. Persidsky, Y., Hill, J., Zhang, M., Dykstra, H., Winfield, M., Reichenbach, N.L., Potula, R., Mukherjee, A., Ramirez, S.H., and Rom, S. (2016). Dysfunction of brain pericytes in chronic neuroinflammation. J Cereb Blood Flow Metab 36, 794–807. 10.1177/0271678X15606149.

33. Bertrand, L., Cho, H.J., and Toborek, M. (2019). Blood-brain barrier pericytes as a target for HIV-1 infection. Brain 142, 502–511. 10.1093/brain/awy339.

34. Bohannon, D.G., Ko, A., Filipowicz, A.R., Kuroda, M.J., and Kim, W.K. (2019). Dysregulation of sonic hedgehog pathway and pericytes in the brain after lentiviral infection. J Neuroinflammation 16, 86. 10.1186/s12974-019-1463-y.

35. Torices, S., Roberts, S.A., Park, M., Malhotra, A., and Toborek, M. (2020). Occludin, caveolin-1, and Alix form a multi-protein complex and regulate HIV-1 infection of brain pericytes. FASEB J 34, 16319–16332. 10.1096/fj.202001562R.

36. Cho, H.J., Kuo, A.M., Bertrand, L., and Toborek, M. (2017). HIV Alters Gap Junction-Mediated Intercellular Communication in Human Brain Pericytes. Frontiers in molecular neuroscience 10, 410. 10.3389/fnmol.2017.00410.

37. Furuse, M., Hirase, T., Itoh, M., Nagafuchi, A., Yonemura, S., Tsukita, S., and Tsukita, S. (1993). Occludin: a novel integral membrane protein localizing at tight junctions. The Journal of cell biology 123, 1777–1788. 10.1083/jcb.123.6.1777.

38. Cong, X., and Kong, W. (2020). Endothelial tight junctions and their regulatory signaling pathways in vascular homeostasis and disease. Cell Signal 66, 109485. 10.1016/j.cellsig.2019.109485.

39. Feldman, G.J., Mullin, J.M., and Ryan, M.P. (2005). Occludin: structure, function and regulation. Adv Drug Deliv Rev 57, 883–917. 10.1016/j.addr.2005.01.009.

40. Cummins, P.M. (2012). Occludin: one protein, many forms. Mol Cell Biol 32, 242–250. 10.1128/MCB.06029-11.

41. Traweger, A., Fang, D., Liu, Y.C., Stelzhammer, W., Krizbai, I.A., Fresser, F., Bauer, H.C., and Bauer, H. (2002). The tight junction-specific protein occludin is a functional target of the E3 ubiquitin-protein ligase itch. J Biol Chem 277, 10201–10208. 10.1074/jbc.M111384200.

42. Choi, Y.B., Shembade, N., Parvatiyar, K., Balachandran, S., and Harhaj, E.W. (2017). TAX1BP1 Restrains Virus-Induced Apoptosis by Facilitating Itch-Mediated Degradation of the Mitochondrial Adaptor MAVS. Mol Cell Biol 37. 10.1128/MCB.00422-16.

43. Castro, V., Skowronska, M., Lombardi, J., He, J., Seth, N., Velichkovska, M., and Toborek, M. (2018). Occludin regulates glucose uptake and ATP production in pericytes by influencing AMP-activated protein kinase activity. J Cereb Blood Flow Metab 38, 317–332. 10.1177/0271678X17720816.

44. Sugiyama, S., Sasaki, T., Tanaka, H., Yan, H., Ikegami, T., Kanki, H., Nishiyama, K., Beck, G., Gon, Y., Okazaki, S., et al. (2023). The tight junction protein occludin modulates blood-brain barrier integrity and neurological function after ischemic stroke in mice. Scientific reports 13, 2892. 10.1038/s41598-023-29894-1.

45. Qi, X., Tang, Z., Shao, X., Wang, Z., Li, M., Zhang, X., He, L., Wang, J., and Yu, X. (2023). Ramelteon improves blood-brain barrier of focal cerebral ischemia rats to prevent post-stroke depression via upregulating occludin. Behavioural brain research 449, 114472. 10.1016/j.bbr.2023.114472.

46. Kim, K.A., Kim, D., Kim, J.H., Shin, Y.J., Kim, E.S., Akram, M., Kim, E.H., Majid, A., Baek, S.H., and Bae, O.N. (2020). Autophagy-mediated occludin degradation contributes to blood-brain barrier disruption during ischemia in bEnd.3 brain endothelial cells and rat ischemic stroke models. Fluids and barriers of the CNS 17, 21. 10.1186/s12987-020-00182-8.

47. Goncalves, A., Su, E.J., Muthusamy, A., Zeitelhofer, M., Torrente, D., Nilsson, I., Protzmann, J., Fredriksson, L., Eriksson, U., Antonetti, D.A., and Lawrence, D.A. (2022). Thrombolytic tPA-induced hemorrhagic transformation of ischemic stroke is mediated by PKCbeta phosphorylation of occludin. Blood 140, 388–400. 10.1182/blood.2021014958.

48. Zhang, Y., Li, X., Qiao, S., Yang, D., Li, Z., Xu, J., Li, W., Su, L., and Liu, W. (2021). Occludin degradation makes brain microvascular endothelial cells more vulnerable to reperfusion injury in vitro. Journal of neurochemistry 156, 352–366. 10.1111/jnc.15102.

49. Costea, L., Meszaros, A., Bauer, H., Bauer, H.C., Traweger, A., Wilhelm, I., Farkas, A.E., and Krizbai, I.A. (2019). The Blood-Brain Barrier and Its Intercellular Junctions in Age-Related Brain Disorders. International journal of molecular sciences 20. 10.3390/ijms20215472.

50. Sweeney, M.D., Sagare, A.P., and Zlokovic, B.V. (2018). Blood-brain barrier breakdown in Alzheimer disease and other neurodegenerative disorders. Nature reviews. Neurology 14, 133–150. 10.1038/nrneurol.2017.188.

51. Li, X., Zhang, Y., Chang, J., Zhang, C., Li, L., Dai, Y., Yang, H., and Wang, Y. (2023). Mfsd2a attenuated hypoxic-ischemic brain damage via protection of the blood-brain barrier in mfat-1 transgenic mice. Cellular and molecular life sciences : CMLS 80, 71. 10.1007/s00018-023-04716-9.

52. Kealy, J., Greene, C., and Campbell, M. (2020). Blood-brain barrier regulation in psychiatric disorders. Neuroscience letters 726, 133664. 10.1016/j.neulet.2018.06.033.

53. Bilgic, A., Ferahkaya, H., Karagoz, H., Kilinc, I., and Energin, V.M. (2023). Serum claudin-5, claudin-11, occludin, vinculin, paxillin, and beta-catenin levels in preschool children with autism spectrum disorder. Nord J Psychiatry 77, 506–511. 10.1080/08039488.2023.2168055.

54. Zengil, S., and Laloglu, E. (2023). Evaluation of Serum Zonulin and Occludin Levels in Bipolar Disorder. Psychiatry investigation 20, 382–389. 10.30773/pi.2022.0234.

55. Leda, A.R., Dygert, L., Bertrand, L., and Toborek, M. (2017). Mouse Microsurgery Infusion Technique for Targeted Substance Delivery into the CNS via the Internal Carotid Artery. Journal of visualized experiments : JoVE. 10.3791/54804.

56. Cuomo, O., Pignataro, G., Gala, R., Scorziello, A., Gravino, E., Piazza, O., Tufano, R., Di Renzo, G., and Annunziato, L. (2007). Antithrombin reduces ischemic volume, ameliorates neurologic deficits, and prolongs animal survival in both transient and permanent focal ischemia. Stroke 38, 3272–3279. 10.1161/STROKEAHA.107.488486.

57. Nahavandi-Parizi, P., Kariminik, A., and Montazeri, M. (2023). Retinoic acid-inducible gene 1 (RIG-1) and IFN-beta promoter stimulator-1 (IPS-1) significantly down-regulated in the severe coronavirus disease 2019 (COVID-19). Molecular biology reports 50, 907–911. 10.1007/s11033-022-07981-2.

58. Peng, Y., Xu, R., and Zheng, X. (2014). HSCARG negatively regulates the cellular antiviral RIG-I like receptor signaling pathway by inhibiting TRAF3 ubiquitination via recruiting OTUB1. PLoS pathogens 10, e1004041. 10.1371/journal.ppat.1004041.

59. Abate, M., Festa, A., Falco, M., Lombardi, A., Luce, A., Grimaldi, A., Zappavigna, S., Sperlongano, P., Irace, C., Caraglia, M., and Misso, G. (2020). Mitochondria as playmakers of apoptosis, autophagy and senescence. Seminars in cell & developmental biology 98, 139–153. 10.1016/j.semcdb.2019.05.022.

60. Wilhelm, L.P., and Ganley, I.G. (2023). Mitochondria and peroxisomes: partners in autophagy. Autophagy 19, 2162–2163. 10.1080/15548627.2022.2155368.

61. Vringer, E., and Tait, S.W.G. (2023). Mitochondria and cell death-associated inflammation. Cell death and differentiation 30, 304–312. 10.1038/s41418-022-01094-w.

62. Hu, Y., Wen, Q., Cai, Y., Liu, Y., Ma, W., Li, Q., Song, F., Guo, Y., Zhu, L., Ge, J., et al. (2023). Alantolactone induces concurrent apoptosis and GSDME-dependent pyroptosis of anaplastic thyroid cancer through ROS mitochondria-dependent caspase pathway. Phytomedicine : international journal of phytotherapy and phytopharmacology 108, 154528. 10.1016/j.phymed.2022.154528.

63. Sun, X., Sun, L., Zhao, Y., Li, Y., Lin, W., Chen, D., and Sun, Q. (2016). MAVS maintains mitochondrial homeostasis via autophagy. Cell Discov 2, 16024. 10.1038/celldisc.2016.24.

64. Patergnani, S., Marchi, S., Rimessi, A., Bonora, M., Giorgi, C., Mehta, K.D., and Pinton, P. (2013). PRKCB/protein kinase C, beta and the mitochondrial axis as key regulators of autophagy. Autophagy 9, 1367–1385. 10.4161/auto.25239.

65. Hu, J., Liu, J., Chen, S., Zhang, C., Shen, L., Yao, K., and Yu, Y. (2023). Thioredoxin-1 regulates the autophagy induced by oxidative stress through LC3-II in human lens epithelial cells. Clinical and experimental pharmacology & physiology 50, 476–485. 10.1111/1440-1681.13764.

66. Tanida, I., Ueno, T., and Kominami, E. (2008). LC3 and Autophagy. Methods in molecular biology 445, 77–88. 10.1007/978-1-59745-157-4_4.

67. Pankiv, S., Clausen, T.H., Lamark, T., Brech, A., Bruun, J.A., Outzen, H., Overvatn, A., Bjorkoy, G., and Johansen, T. (2007). p62/SQSTM1 binds directly to Atg8/LC3 to facilitate degradation of ubiquitinated protein aggregates by autophagy. J Biol Chem 282, 24131–24145. 10.1074/jbc.M702824200.

68. Ma, K., Wu, H., Li, P., and Li, B. (2018). LC3-II may mediate ATR-induced mitophagy in dopaminergic neurons through SQSTM1/p62 pathway. Acta biochimica et biophysica Sinica 50, 1047–1061. 10.1093/abbs/gmy091.

69. Viret, C., Rozieres, A., and Faure, M. (2018). Novel Insights into NDP52 Autophagy Receptor Functioning. Trends Cell Biol 28, 255–257. 10.1016/j.tcb.2018.01.003.

70. Abu Shelbayeh, O., Arroum, T., Morris, S., and Busch, K.B. (2023). PGC-1alpha Is a Master Regulator of Mitochondrial Lifecycle and ROS Stress Response. Antioxidants 12. 10.3390/antiox12051075.

71. Li, Y., Hei, H., Zhang, S., Gong, W., Liu, Y., and Qin, J. (2023). PGC-1alpha participates in tumor chemoresistance by regulating glucose metabolism and mitochondrial function. Molecular and cellular biochemistry 478, 47–57. 10.1007/s11010-022-04477-2.

72. Fernandez-Trujillo, M.A., Garcia-Rosado, E., Alonso, M.C., Castro, D., Alvarez, M.C., and Bejar, J. (2013). Mx1, Mx2 and Mx3 proteins from the gilthead seabream (Sparus aurata) show in vitro antiviral activity against RNA and DNA viruses. Molecular immunology 56, 630–636. 10.1016/j.molimm.2013.06.018.

73. Kim, W.K., Avarez, X., and Williams, K. (2005). The role of monocytes and perivascular macrophages in HIV and SIV neuropathogenesis: information from non-human primate models. Neurotoxicity research 8, 107–115. 10.1007/BF03033823.

74. Chen, N.C., Partridge, A.T., Sell, C., Torres, C., and Martin-Garcia, J. (2017). Fate of microglia during HIV-1 infection: From activation to senescence? Glia 65, 431–446. 10.1002/glia.23081.

75. Bai, R., Song, C., Lv, S., Chang, L., Hua, W., Weng, W., Wu, H., and Dai, L. (2023). Role of microglia in HIV-1 infection. AIDS research and therapy 20, 16. 10.1186/s12981-023-00511-5.

76. Wilson, K.M., and He, J.J. (2023). HIV Nef Expression Down-modulated GFAP Expression and Altered Glutamate Uptake and Release and Proliferation in Astrocytes. Aging and disease 14, 152–169. 10.14336/AD.2022.0712.

77. Ziegler, C.G.K., Allon, S.J., Nyquist, S.K., Mbano, I.M., Miao, V.N., Tzouanas, C.N., Cao, Y., Yousif, A.S., Bals, J., Hauser, B.M., et al. (2020). SARS-CoV-2 Receptor ACE2 Is an Interferon-Stimulated Gene in Human Airway Epithelial Cells and Is Detected in Specific Cell Subsets across Tissues. Cell 181, 1016–1035 e1019. 10.1016/j.cell.2020.04.035.

78. Levy, D.E., and Garcia-Sastre, A. (2001). The virus battles: IFN induction of the antiviral state and mechanisms of viral evasion. Cytokine Growth Factor Rev 12, 143–156. 10.1016/s1359-6101(00)00027-7.

79. Verhelst, J., Hulpiau, P., and Saelens, X. (2013). Mx proteins: antiviral gatekeepers that restrain the uninvited. Microbiol Mol Biol Rev 77, 551–566. 10.1128/MMBR.00024-13.

80. Xu, X.Q., Liu, Y., Zhang, B., Liu, H., Shao, D.D., Liu, J.B., Wang, X., Zhou, L.N., Hu, W.H., and Ho, W.Z. (2019). IL-22 suppresses HSV-2 replication in human cervical epithelial cells. Cytokine 123, 154776. 10.1016/j.cyto.2019.154776.

81. Betancor, G., Dicks, M.D.J., Jimenez-Guardeno, J.M., Ali, N.H., Apolonia, L., and Malim, M.H. (2019). The GTPase Domain of MX2 Interacts with the HIV-1 Capsid, Enabling Its Short Isoform to Moderate Antiviral Restriction. Cell Rep 29, 1923–1933 e1923. 10.1016/j.celrep.2019.10.009.

82. Dicks, M.D.J., Betancor, G., Jimenez-Guardeno, J.M., Pessel-Vivares, L., Apolonia, L., Goujon, C., and Malim, M.H. (2018). Multiple components of the nuclear pore complex interact with the amino-terminus of MX2 to facilitate HIV-1 restriction. PLoS Pathog 14, e1007408. 10.1371/journal.ppat.1007408.

83. Spitaels, J., Van Hoecke, L., Roose, K., Kochs, G., and Saelens, X. (2019). Mx1 in Hematopoietic Cells Protects against Thogoto Virus Infection. J Virol 93. 10.1128/JVI.00193-19.

84. Li, N., Zhang, L., Chen, L., Feng, W., Xu, Y., Chen, F., Liu, X., Chen, Z., and Liu, W. (2012). MxA inhibits hepatitis B virus replication by interaction with hepatitis B core antigen. Hepatology 56, 803–811. 10.1002/hep.25608.

85. Haller, O., and Kochs, G. (2019). Mx genes: host determinants controlling influenza virus infection and trans-species transmission. Hum Genet. 10.1007/s00439-019-02092-8.

86. Staeheli, P., and Haller, O. (2018). Human MX2/MxB: a Potent Interferon-Induced Postentry Inhibitor of Herpesviruses and HIV-1. J Virol 92. 10.1128/JVI.00709-18.

87. Bertrand, L., Meroth, F., Tournebize, M., Leda, A.R., Sun, E., and Toborek, M. (2019). Targeting the HIV-infected brain to improve ischemic stroke outcome. Nat Commun 10, 2009. 10.1038/s41467-019-10046-x.

88. Qin, Z., Fang, X., Sun, W., Ma, Z., Dai, T., Wang, S., Zong, Z., Huang, H., Ru, H., Lu, H., et al. (2022). Deactylation by SIRT1 enables liquid-liquid phase separation of IRF3/IRF7 in innate antiviral immunity. Nature immunology 23, 1193–1207. 10.1038/s41590-022-01269-0.

89. Ning, S., Pagano, J.S., and Barber, G.N. (2011). IRF7: activation, regulation, modification and function. Genes Immun 12, 399–414. 10.1038/gene.2011.21.

90. Mogensen, T.H. (2018). IRF and STAT Transcription Factors -From Basic Biology to Roles in Infection, Protective Immunity, and Primary Immunodeficiencies. Front Immunol 9, 3047. 10.3389/fimmu.2018.03047.

91. Chan, D.C. (2020). Mitochondrial Dynamics and Its Involvement in Disease. Annual review of pathology 15, 235–259. 10.1146/annurev-pathmechdis-012419-032711.

92. Loson, O.C., Song, Z., Chen, H., and Chan, D.C. (2013). Fis1, Mff, MiD49, and MiD51 mediate Drp1 recruitment in mitochondrial fission. Molecular biology of the cell 24, 659–667. 10.1091/mbc.E12-10-0721.

93. Saha, K., Subramenium Ganapathy, A., Wang, A., Michael Morris, N., Suchanec, E., Ding, W., Yochum, G., Koltun, W., Nighot, M., Ma, T., and Nighot, P. (2023). Autophagy Reduces the Degradation and Promotes Membrane Localization of Occludin to Enhance the Intestinal Epithelial Tight Junction Barrier against Paracellular Macromolecule Flux. Journal of Crohn’s & colitis 17, 433–449. 10.1093/ecco-jcc/jjac148.

94. Nong, H., Yuan, H., Lin, Y., Chen, S., Li, Y., Luo, Z., Yang, W., Zhang, T., and Chen, Y. (2023). IL-22 promotes occludin expression by activating autophagy and treats ulcerative colitis. Molecular and cellular biochemistry. 10.1007/s11010-023-04806-z.

95. Li, Y., Zhang, P., Zhang, J., Bao, W., Li, J., Wei, Y., Ni, J., and Gong, K. (2022). Role of Autophagy Inducers and Inhibitors in Intestinal Barrier Injury Induced by Intestinal Ischemia-Reperfusion (I/R). Journal of immunology research 2022, 9822157. 10.1155/2022/9822157.

96. Rachow, S., Zorn-Kruppa, M., Ohnemus, U., Kirschner, N., Vidal-y-Sy, S., von den Driesch, P., Bornchen, C., Eberle, J., Mildner, M., Vettorazzi, E., et al. (2013). Occludin is involved in adhesion, apoptosis, differentiation and Ca2+-homeostasis of human keratinocytes: implications for tumorigenesis. PloS one 8, e55116. 10.1371/journal.pone.0055116.

97. Kuo, W.T., Shen, L., Zuo, L., Shashikanth, N., Ong, M., Wu, L., Zha, J., Edelblum, K.L., Wang, Y., Wang, Y., et al. (2019). Inflammation-induced Occludin Downregulation Limits Epithelial Apoptosis by Suppressing Caspase-3 Expression. Gastroenterology 157, 1323–1337. 10.1053/j.gastro.2019.07.058.

98. Kitajiri, S., Katsuno, T., Sasaki, H., Ito, J., Furuse, M., and Tsukita, S. (2014). Deafness in occludin-deficient mice with dislocation of tricellulin and progressive apoptosis of the hair cells. Biology open 3, 759–766. 10.1242/bio.20147799.

99. Gu, J.M., Lim, S.O., Park, Y.M., and Jung, G. (2008). A novel splice variant of occludin deleted in exon 9 and its role in cell apoptosis and invasion. The FEBS journal 275, 3145–3156. 10.1111/j.1742-4658.2008.06467.x.

100. Osanai, M., Murata, M., Nishikiori, N., Chiba, H., Kojima, T., and Sawada, N. (2006). Epigenetic silencing of occludin promotes tumorigenic and metastatic properties of cancer cells via modulations of unique sets of apoptosis-associated genes. Cancer research 66, 9125–9133. 10.1158/0008-5472.CAN-06-1864.

101. Potash, M.J., Chao, W., Bentsman, G., Paris, N., Saini, M., Nitkiewicz, J., Belem, P., Sharer, L., Brooks, A.I., and Volsky, D.J. (2005). A mouse model for study of systemic HIV-1 infection, antiviral immune responses, and neuroinvasiveness. Proc Natl Acad Sci U S A 102, 3760–3765. 10.1073/pnas.0500649102.

102. Kramer, M., Dang, J., Baertling, F., Denecke, B., Clarner, T., Kirsch, C., Beyer, C., and Kipp, M. (2010). TTC staining of damaged brain areas after MCA occlusion in the rat does not constrict quantitative gene and protein analyses. Journal of neuroscience methods 187, 84–89. 10.1016/j.jneumeth.2009.12.020.

103. Love, M.I., Huber, W., and Anders, S. (2014). Moderated estimation of fold change and dispersion for RNA-seq data with DESeq2. Genome Biol 15, 550. 10.1186/s13059-014-0550-8.

104. McArthur, J.C., Nance-Sproson, T.E., Griffin, D.E., Hoover, D., Selnes, O.A., Miller, E.N., Margolick, J.B., Cohen, B.A., Farzadegan, H., and Saah, A. (1992). The diagnostic utility of elevation in cerebrospinal fluid beta 2-microglobulin in HIV-1 dementia. Multicenter AIDS Cohort Study. Neurology 42, 1707–1712. 10.1212/wnl.42.9.1707.

105. Shannon, P., Markiel, A., Ozier, O., Baliga, N.S., Wang, J.T., Ramage, D., Amin, N., Schwikowski, B., and Ideker, T. (2003). Cytoscape: a software environment for integrated models of biomolecular interaction networks. Genome Res 13, 2498–2504. 10.1101/gr.1239303.

106. Merico, D., Isserlin, R., Stueker, O., Emili, A., and Bader, G.D. (2010). Enrichment map: a network-based method for gene-set enrichment visualization and interpretation. PLoS One 5, e13984. 10.1371/journal.pone.0013984.

107. Reimand, J., Isserlin, R., Voisin, V., Kucera, M., Tannus-Lopes, C., Rostamianfar, A., Wadi, L., Meyer, M., Wong, J., Xu, C., et al. (2019). Pathway enrichment analysis and visualization of omics data using g:Profiler, GSEA, Cytoscape and EnrichmentMap. Nat Protoc 14, 482–517. 10.1038/s41596-018-0103-9.

108. Walter, W., Sanchez-Cabo, F., and Ricote, M. (2015). GOplot: an R package for visually combining expression data with functional analysis. Bioinformatics 31, 2912–2914. 10.1093/bioinformatics/btv300.

109. Valente, A.J., Maddalena, L.A., Robb, E.L., Moradi, F., and Stuart, J.A. (2017). A simple ImageJ macro tool for analyzing mitochondrial network morphology in mammalian cell culture. Acta histochemica 119, 315–326. 10.1016/j.acthis.2017.03.001.

